# ACVR1^R206H^ increases osteogenic/ECM gene expression and impairs myofiber formation in human skeletal muscle stem cells

**DOI:** 10.1101/2021.01.18.427082

**Authors:** Emilie Barruet, Steven M. Garcia, Jake Wu, Blanca M. Morales, Stanley Tamaki, Tania Moody, Jason H. Pomerantz, Edward C. Hsiao

**Affiliations:** Division of Endocrinology and Metabolism, Department of Medicine, and the Institute for Human Genetics, University of California, San Francisco, CA 94143, U.S.A.; Departments of Surgery and Orofacial Sciences, Division of Plastic and Reconstructive Surgery, Program in Craniofacial Biology, Eli and Edythe Broad Center of Regeneration Medicine, University of California San Francisco, 94143, USA; Department of Surgery, Division of Plastic and Reconstructive Surgery, University of California, San Francisco, 94143, USA

## Abstract

Abnormalities in skeletal muscle repair lead to poor function and complications such as scarring or heterotopic ossification (HO). Here, we use fibrodysplasia ossificans progressiva (FOP), a disease of progressive HO caused by ACVR1^R206H^ (Activin receptor type-1 receptor) mutation, to elucidate how ACVR1 affects skeletal muscle repair. Rare and unique primary FOP human muscle stem cells (Hu-MuSCs) isolated from cadaveric skeletal muscle demonstrated increased ECM marker expression, and showed skeletal muscle-specific impaired engraftment and regeneration ability. Human induced pluripotent stem cell (iPSC)-derived muscle stem/progenitor cells (iMPCs) Single cell transcriptome analyses from FOP also revealed unusually increased ECM and osteogenic marker expression compared to control iMPCs. These results show that iMPCs can recapitulate many aspects of Hu-MuSCs for detailed in vitro study, that ACVR1 is a key regulator of Hu-MuSC function and skeletal muscle repair; and that ACVR1 activation in iMPCs or Hu-MuSCs contributes to HO by changing the local tissue environment.

## Introduction

Human diseases of skeletal muscle are major medical problems. Aberrant repair after muscle injury results in scarring or heterotopic ossification (HO; bone formation in an inappropriate site) which can be devastating. Disrupted signaling of bone morphogenetic proteins (BMPs), originally identified by their ability to induce bone formation when injected into muscle (Urist, 1965), changes muscle homeostasis (Ono et al., 2011) by controlling the proliferation and differentiation of satellite cells (SCs) (Ono et al., 2011; Stantzou et al., 2017). SCs marked by PAX7 (Paired Box 7) (Seale et al., 2000), are a prerequisite for skeletal muscle regeneration (Mauro, 1961) and are thought to be the main human muscle stem cells (Hu-MuSCs) of postnatal skeletal muscle. Upon injury, activated SCs give rise to myoblasts, which form new myofibers or fuse to existing muscle fibers to repair muscle damage (Kuang, Kuroda, Le Grand, & Rudnicki, 2007). A subset of SCs does not differentiate and serves to replenish the SC pool. However, how the BMP pathway regulates muscle repair or SC function in conditions of HO remains unclear and is a major knowledge gap.

Recent protocols to create Hu-MuSCs-like cells from human induced pluripotent stem cells (hiPSCs) (Takahashi et al., 2007) could generate skeletal myogenic lineage cells, but with limited muscle regenerative capacity (Borchin, Chen, & Barberi, 2013; Magli et al., 2017; Shelton et al., 2014; van der Wal et al., 2018; Xi et al., 2017). PAX7 cell engraftment capability was often not reported or showed limited success. Transgene-induced *PAX7* or *PAX3* expression in hiPSCs can increase engraftment of PAX7^+^ cells and contribution to the SC pool (Al Tanoury et al., 2020; Wu et al., 2018); however, these engineered cells may not reflect physiology due to the genetic manipulation of these master transcription factors.

BMP signaling is one pathway that could be manipulated to make Hu-MuSCs-like cells from hiPSCs (iMPCs) (Chal et al., 2016). Although inhibiting the BMP pathway is an important step that promotes iMPC formation, the BMP pathway is also critical for maintaining *PAX7* expression in primary MuSCs and for preventing commitment to myogenic differentiation (Friedrichs et al., 2011). Abrogation of BMP signaling in SCs slowed myofiber growth (Stantzou et al., 2017), and increased BMP4 levels in Duchenne’ s muscular dystrophy (DMD) can exacerbate the disease (Shi, de Gorter, Hoogaars, t Hoen, & ten Dijke, 2013).

Fibrodysplasia ossificans progressiva (FOP), a congenital disease of abnormal skeletal muscle regeneration and severe HO, provides a unique opportunity to understand how changes in BMP signaling affect Hu-MuSC and iMPC formation and function. The FOP iMPCs and Hu-MuSCs carry the classical ACVR1^R206H^ (c.617>A) mutation (Shore et al., 2006) that causes hyperactivation of the BMP-SMAD signaling (Billings et al., 2008) and aberrant responses to Activin A (Hatsell et al., 2015). Primary FOP Hu-MuSCs can engraft and regenerate injured muscle of NSG mice, but at lower efficiency than Hu-MuSCs from non-FOP donors, and that the source of FOP Hu-MuSCs (biceps vs. diaphragm) appears to impact their engraftment efficiency. A new non-transgenic strategy for creating human iMPCs was developed and applied to multiple iPSC lines from control subjects and subjects with FOP. These iMPCs shared transcriptional similarities with primary Hu-MuSCs, and abnormal activation of the BMP pathway by ACVR1^R206H^ changed the transcription profiles of the FOP iMPCs compared to controls.

## Results

### 1 Lower efficiency engraftment of Primary Human FOP Hu-MuSCs

Muscle tissue samples were obtained from FOP cadavers which allowed the study of markers *in situ* and the isolation of primary human cells with endogenous activated ACVR1 signaling. Hematoxylin and eosin and alcian blue staining of FOP primary muscle samples from two deceased FOP subjects (**Figure 1-Source Data 1**) in a region without HO showed no gross defects. FOP muscle tissue near a HO lesion showed increased ECM proteoglycan components (alcian blue staining) in the interstitial space of the muscle fibers near the HO lesion (**Figure 1A**). Primary Hu-MuSCs carrying the ACVR1^R206H^ activating mutation were isolated from unfixed muscle tissue from two FOP autopsies. Biceps muscle, commonly affected by HO, and diaphragm muscle, one of the rare skeletal muscle sites spared from HO in patients with FOP, were analyzed. PAX7 staining confirmed the presence of SCs in human FOP biceps (**Figure 1B**, top and middle). Interestingly, a small subregion showed Collagen Type 1 expression with embedded satellite cells (**Figure 1B**, bottom) suggesting possible early HO formation despite absence of gross HO.

**Figure 1:**
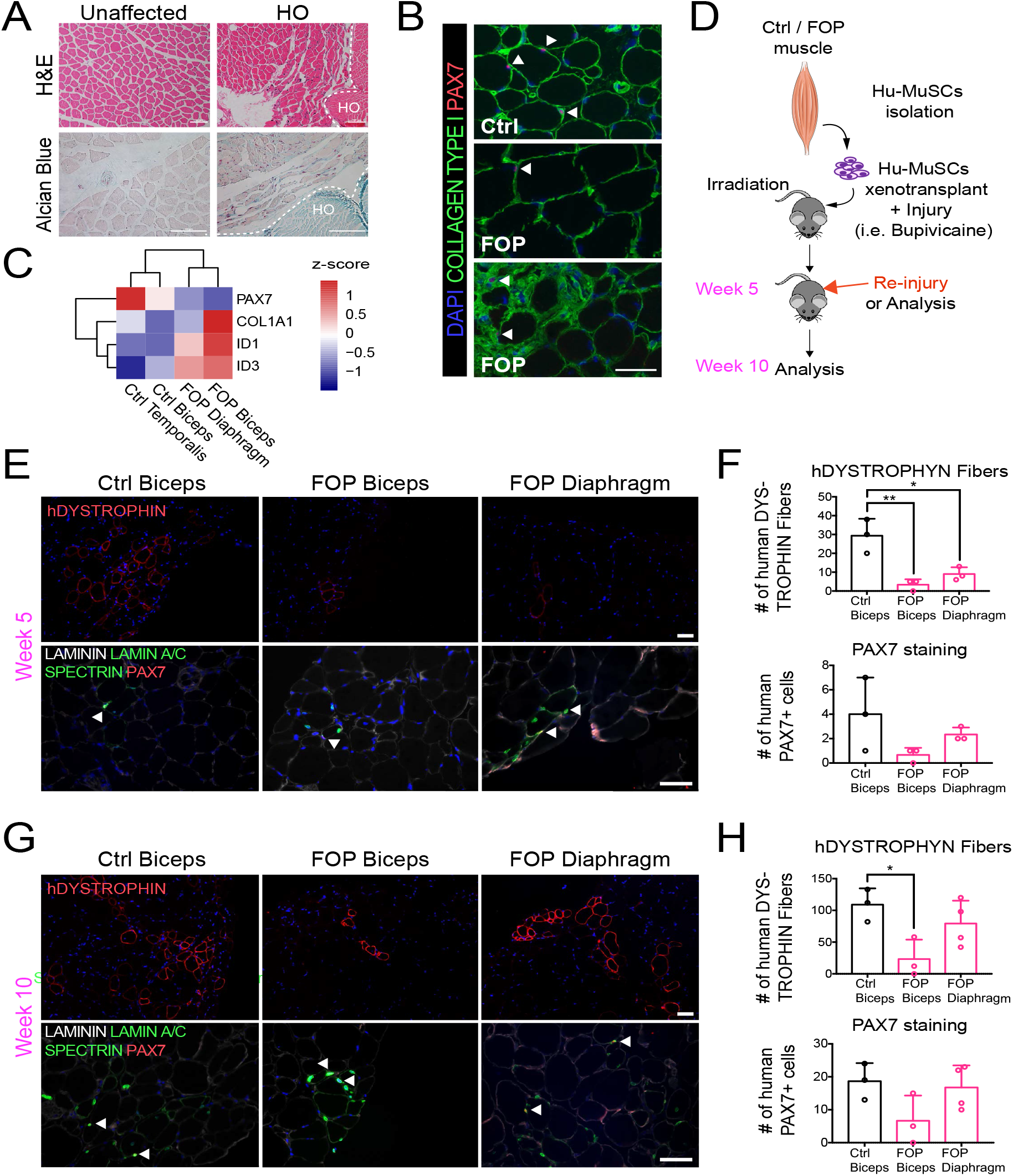
Regenerative properties of primary Hu-MuSCs isolated from FOP patients. **(A)** Hematoxylin and eosin staining of muscle cross sections from FOP subjects, with no heterotopic ossification (left) and with heterotopic ossification, depicted by the white dashed line (right, 100µm scale bar). **(B)** Immunofluorescence staining for PAX7 and COL1 of muscle cross sections from control and FOP subjects. White arrowheads mark SCs, 50µmscale bar. **(C)** Heat map of normalized gene expression of sorted human satellite cells from 2 control subjects and 2 different muscles of 1 FOP subject. **(D)** Schematic and experimental time course. SCs from biceps and diaphragm muscles of a deceased FOP patient were sorted and transplanted into NSG mice. **(E)** Immunofluorescence staining 5 weeks after xenotranplantation. White arrowheads mark SCs, 50 µm scale bar. **(F)** Quantification of human DYSTROPHIN fibers and human PAX7^+^ cells 5 weeks after transplant. **(G)** Mice were re-injured 5 weeks after transplantation with bupivacaine. Immunofluorescence staining was at week 10. White arrows mark SCs, 50µm scale bar. **(H)** Quantification of human DYSTROPHIN fibers and human PAX7^+^ cells after re-injury at week 10. n=1, biological replicates, n≥3 technical replicates. Error bar represent mean and SD. *, p<0.05, **, p<0.01. Muscle specimen details are listed in **Figure 1-Source Data 1**.

Sufficient Hu-MuSCs (**Figure 1-figure supplement 1A**) were sorted for transplant and gene expression analysis (**Figure 1-Source Data 2**). FOP Hu-MuSCs showed lower *PAX7* expression compared to control Hu-MuSCs. *COL1A1* and *ID3* were increased in affected FOP muscle (biceps) (**Figure 1C**). However, *ID1* was increased in both the FOP diaphragm and biceps compared to control muscles (**Figure 1C**). Five weeks after transplantation, biceps and diaphragm FOP Hu-MuSCs had engrafted and formed human fibers (**Figure 1D, E**). However, the number of human DYSTROPHIN-positive fibers was significantly lower with FOP Hu-MuSCs vs. control Hu-MuSCs (**Figure 1F**). The number of engrafted PAX7^+^ cells was qualitatively lower with FOP vs. control Hu-MuSCs (**Figure 1F** and **Figure 1-figure supplement 1B**). Ten weeks after re-injury with bupivacaine at week 5 (**Figure 1D**), the number of human DYSTROPHIN fibers was significantly decreased when FOP biceps Hu-MuSCs were transplanted compared to control Hu-MuSCs, but not when FOP diaphragm Hu-MuSCs were transplanted (**Figure 1G, H**). No differences in the number of human PAX7 cells were identified (**Figure 1G, H**). No radiologic evidence of HO was found in any mice (**Figure 1-figure supplement 1C**).

Thus, primary FOP Hu-MuSCs can engraft and regenerate injured muscle of NSG mice, but at lower efficiency than unaffected MuSCs, and the source of FOP Hu-MuSCs (biceps or diaphragm) may impact MuSC engraftment efficiency.

### 2 Human FOP iPSCs can differentiate into skeletal muscle cells

The rarity of FOP disease and difficulty obtaining human tissue samples from patients with FOP makes it difficult to obtain a reliable source of muscle stem cells. Therefore, we used established and fully characterized control hiPSCs (Wtc11, 1323-2, and BJ2) and ACVR1^R206H^ hiPSCs (F1-1, F2-3, F3-2) lines previously derived from patients with FOP. The control hiPSCs yielded PAX7 and MYOGENIN-expressing cells (**Figure 2A,B** and **Figure 2-figure supplement 1A,B**) as early as day 25 (**Figure 2-figure supplement 1A,B**), expressed DYSTROPHIN (**Figure 2B**), and formed contractile myotubes (**Figure 2C** and **Video1-2**).

**Figure 2:**
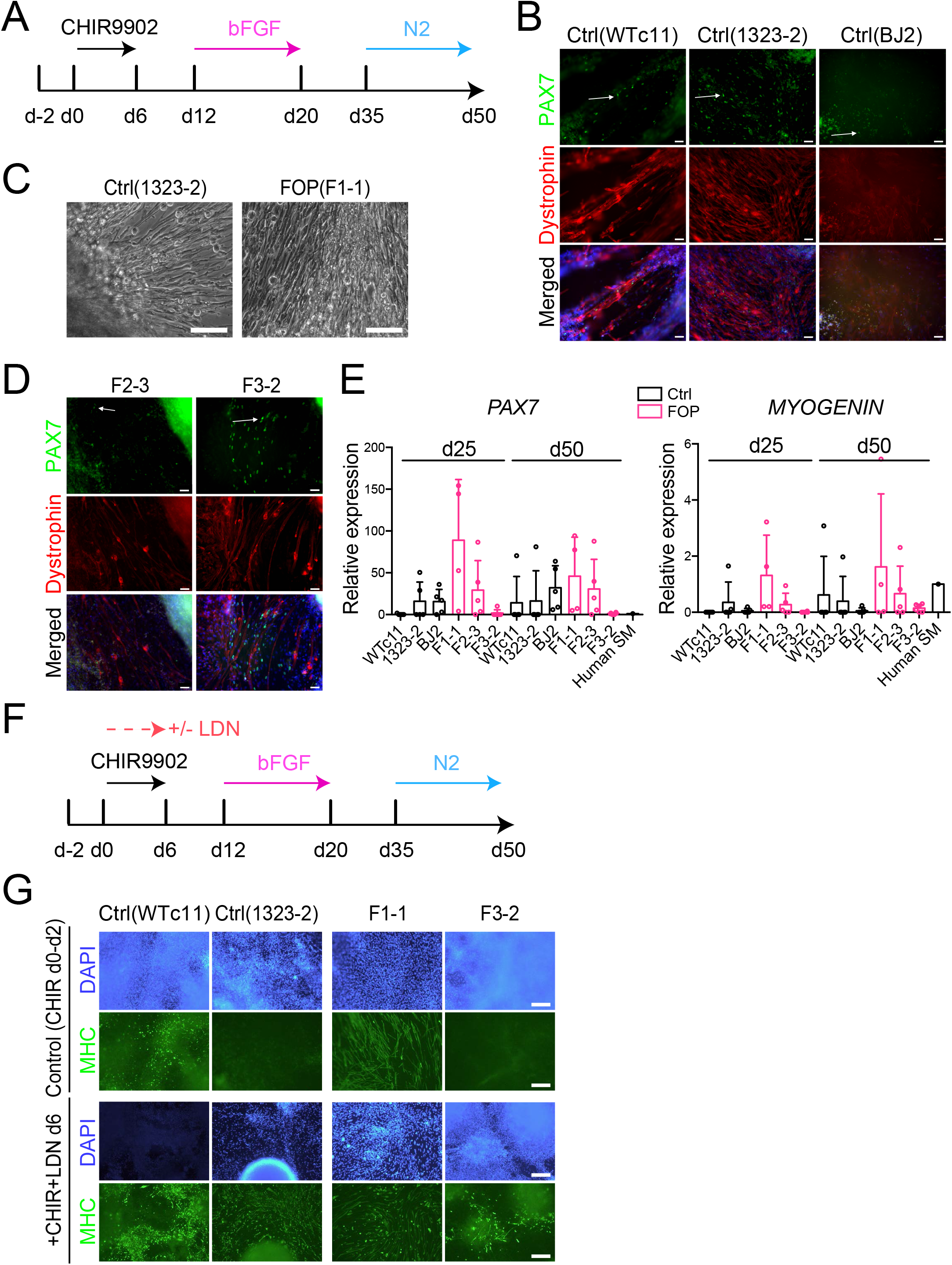
Skeletal muscle differentiation of human control and FOP iPSC lines. **(A)** Differentiation schematic of hiPSCs into skeletal muscle stem cell-like cells (iMPCs). **(B)** Immunofluorescence staining of iMPCs at day 50. White arrows show PAX7 expressing cells, 50µm scale bar. **(C)** iMPCs can form contractile myotubes, 100µm scale bar. **(D)** FOP iMPCs form DYSTROPHIN and PAX7 expressing cells (day 50). White arrows show PAX7 expressing cells, 50µm scale bar. **(E)** *PAX7* and *MYOGENIN* gene expression at day 25 and day 50 of differentiation (n=3 biological replicates and n≥3 technical replicates). Error bars represent mean ± SD. No statistically significant differences were detected. **(F)** Schematic describing the addition of the BMP pathway inhibitor LDN193189 (LDN). **(G)** Representative immunofluorescence staining of 2 control and 2 FOP iMPCs for MHC (Myosin Heavy Chain, green) and nuclear stain DAPI (blue) at day 50 +/- LDN addition, 200µm scale bar

Since BMPs control skeletal muscle differentiation from hiPSCs (Chal et al., 2016; Xi et al., 2017), we investigated if genetic activation of the BMP pathway via ACVR1^R206H^ could alter the myogenic differentiation of hiPSCs (F1-1, F2-3, F3-2) derived from patients with FOP (Matsumoto et al., 2013). All three FOP iPSC lines formed contractile myotubes with cells expressing DYSTROPHIN and PAX7 (**Figure 2C,D and Figure 2-figure supplement 1A,B**). *PAX7, MYOGENIN*, and *DYSTROPHIN* (**Figure 2E** and **Figure 2-figure supplement 1C**) gene expression showed some heterogeneity among the different hiPSC lines but these differences were not statistically significant. Adding a SMAD inhibitor of the BMP pathway (LDN193189) into the differentiation protocol (**Figure 2F**) improved the differentiation of the control hiPSC lines. LDN also improved differentiation of the FOP F3-2 (**Figure 2G**) line, which had shown lower PAX7 and MYOGENIN expression (**Figure 2E**). These findings show that individual hiPSC lines are heterogeneous in differentiation to skeletal muscle lineages, similar to other protocols (Volpato & Webber, 2020), that FOP hiPSCs can form skeletal muscle cells and Hu-MuSC-like cells expressing PAX7 despite upregulation of the BMP pathway by ACVR1^R206H^, and that chemical blockade of the BMP pathway can improve the formation of Hu-MuSC-like cells from the FOP iPSC line that showed the lowest efficiency.

### 3 PAX7 expressing cells can be isolated from myogenic differentiation

To test the regenerative properties of PAX7-expressing MuSCs, we used FACS to purify HNK1^-^ CD45^-^CD31^-^ cells co-expressing CD29, CXCR4, and CD56 markers present on human PAX7^+^ cells (Garcia et al., 2018) (**Figure 3A** and **Figure 3-figure supplement 1A,B**). FACS analysis identified intermediate CD56 cells expressing high *PAX7* and low *MYOGENIN* (**Figure 3B** and **Figure 3-figure supplement 1C**), consistent with Hu-MuSC expression profiles. All HNK1^-^ CD45^-^CD31^-^ CXCR4^+^CD29^+^CD56^dim^ cells formed myotubes expressing MHC (**Figure 3-figure supplement 1D**) when cultured in terminal differentiation media demonstrating isolation of functional iMPCs with satellite cell characteristics from the cultures.

**Figure 3:**
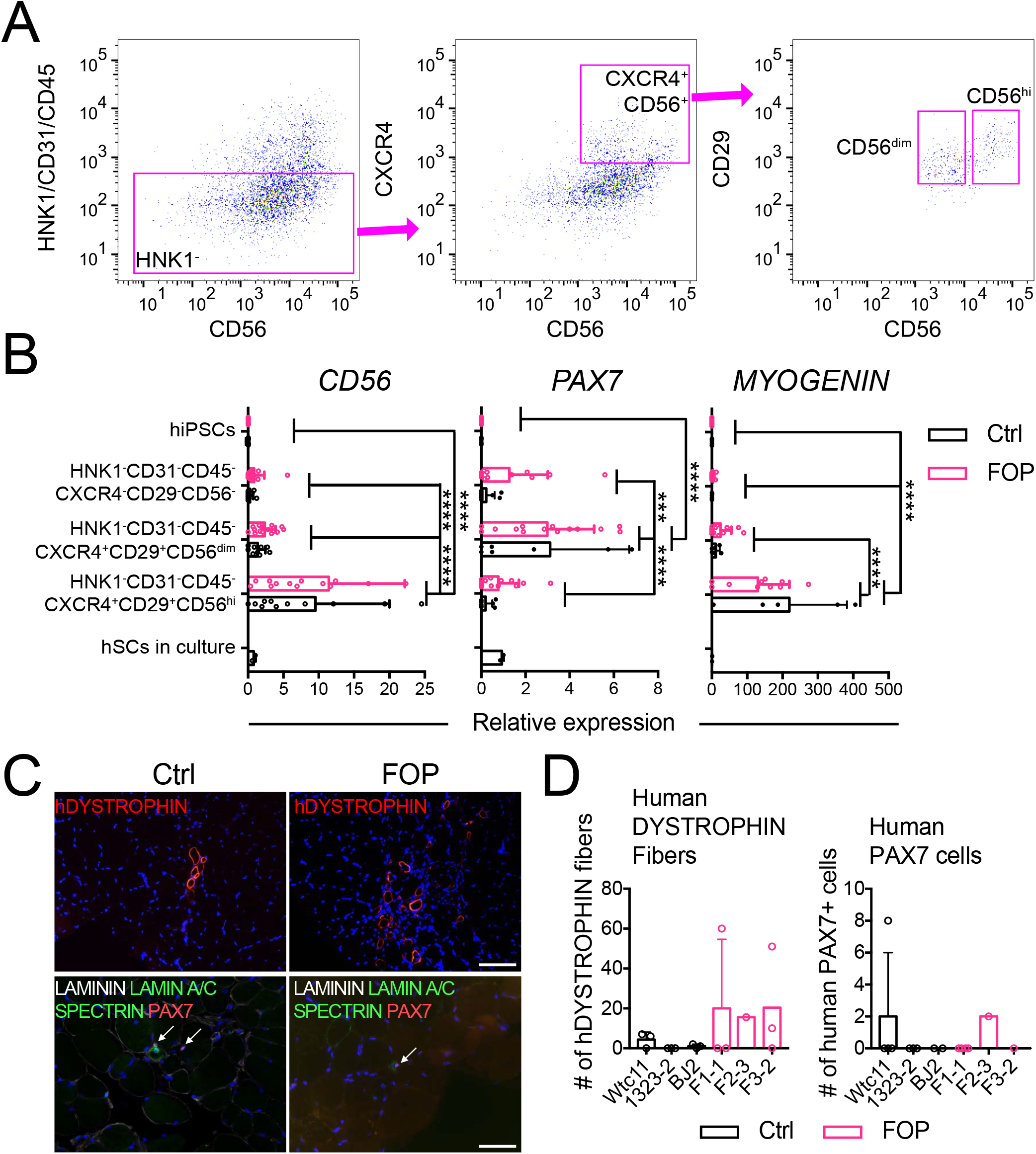
Isolation and transplantation of PAX7-expressing cells from iPSC muscle differentiation culture. hiPSCs were differentiated into iMPCs until day 50 and sorted via flow cytometry. **(A)** Gating strategy. **(B)** Myogenic gene expression of CD56^dim^ and CD56^hi^ cells (n=3 biological and technical replicates, \*\*\**p* < 0.001, \*\*\*\**p* < 0.0001 by two-way ANOVA test). **(C)** Representative human DYSTROPHIN (top, 200 µm scale bar), and total LAMININ, human LAMIN A/C, human SPECTRIN, and PAX7 (bottom, 100 µm scale bar) immunohistochemistry of NSG mice anterior tibialis, where sorted iMPCs were transplanted. White arrows show engrafted hiPSC-derived muscle stem cells. **(D)** Quantification of human DYSTROPHIN fibers and human PAX7 cells at week 5 after transplant (n=3 biological replicates).

### 4 Isolated iMPCs can regenerate injured mouse muscle and form human fibers

To assess iMPC regenerative capacity *in vivo*, we injected 1,000-10,000 iMPCs derived from control or FOP hiPSCs (**Figure 3-Source Data 1**) into the tibialis anterior (TA) muscle of whole-body irradiated NSG immunocompromised mice (to hinder endogenous satellite cells) previously injured with bupivacaine (Garcia, Tamaki, Xu, & Pomerantz, 2017). The bupivicaine step induces myofiber injury and promotes engraftment of donor SCs. New fibers expressing human DYSTROPHIN and PAX7 cells were found after 5 weeks, showing that iMPCs could engraft and promote muscle regeneration (**Figure 3C**). However, the number of human fibers and human PAX7^+^ cells remained low (**Figure 3D**) compared to primary Hu-MuSCs. While some iMPC transplants yielded up to 60 new human fibers, some did not yield any human fibers. By comparison, 2,000 primary non-FOP Hu-MuSCs resulted in an average of 155 human fibers based on prior assessments using the same assay (Garcia et al., 2018). No significant differences between control and FOP iMPCs were identified, though some individual FOP samples showed higher engraftment (**Figure 3D**).

These results showed that iMPCs can engraft into a muscle injury site in mice, but engraftment efficiency may be lower than primary Hu-MuSCs or be the result of differences in experimental conditions. Also, ACVR1^R206H^ did not significantly impact muscle fiber regeneration in this assay.

### 5 Transcriptional profiling of iMPCs

The lower engraftment of iMPCs compared to primary Hu-MuSCs suggested that the FACS-purified population was still heterogeneous or that iMPCs do not fully recapitulate adult primary Hu-MuSCs. Single cell RNA sequencing (scRNAseq) from control (1323-2) and FOP (F3-2) iMPCs (the lines were selected based on their lower intra-line variability, **Figure 2**) were analyzed (**Figure 4A; Figure 4-Source Data 1)**. Cell populations for both samples were defined by the dimension reduction technique of uniform manifold approximation and projection (UMAP) (Becht et al., 2018) and unsupervised clustering with Seurat v3 package (Stuart et al., 2019) **Figure 4B**). Both control (**Figure 4-figure supplement 1A**) and FOP (**Figure 4-figure supplement 1B**) samples had clusters expressing myogenic genes (*PAX7* and *MYOD*); mesenchymal genes (*PDGFRA*); and neuronal genes (*SOX2*). Merged analysis to allow direct comparison identified 13 distinct clusters (**Figure 4C** and **Figure 4-figure supplement 1C**). Clusters 0-2, 4-9, and 12 consisted of cells expressing neuronal stem/progenitor cell genes such as *EFNB3, SOX2*, and *ASCL1* (**Figure 4D,E** and **Figure 4-figure supplement 1D,E**). Mesenchymal genes (*PDGFRA, ASPN*, and *COL1A1*) were expressed in cluster 8 (**Figure 4D,E** and **Figure 4-figure supplement 1D,E**). Cells expressing muscle markers (*PAX7, MYF5, MYOD* and *MYOG* (*MYOGENIN*)) were found in clusters 3, 10, and 11 (**Figure 4D,E** and **Figure 4-figure supplement 1D,E**). The frequency of muscle cells (clusters 3, 10, 11) was higher in FOP (14%, 2.5%, and 1.7%) compared to controls (4.2%, 0.5%, and 0.5%) (**Figure 4F**) suggesting that FOP hiPSCs may be more efficient at making muscle progenitor cells. Thus, FACS-purified iMPC cultures contain muscle stem/progenitor cells, but other cell types such as mesenchymal, neuronal progenitor cells, and myoblasts persist in the culture. Furthermore, the frequency of iMPCs appears to be higher in FOP vs. control cell cultures.

**Figure 4:**
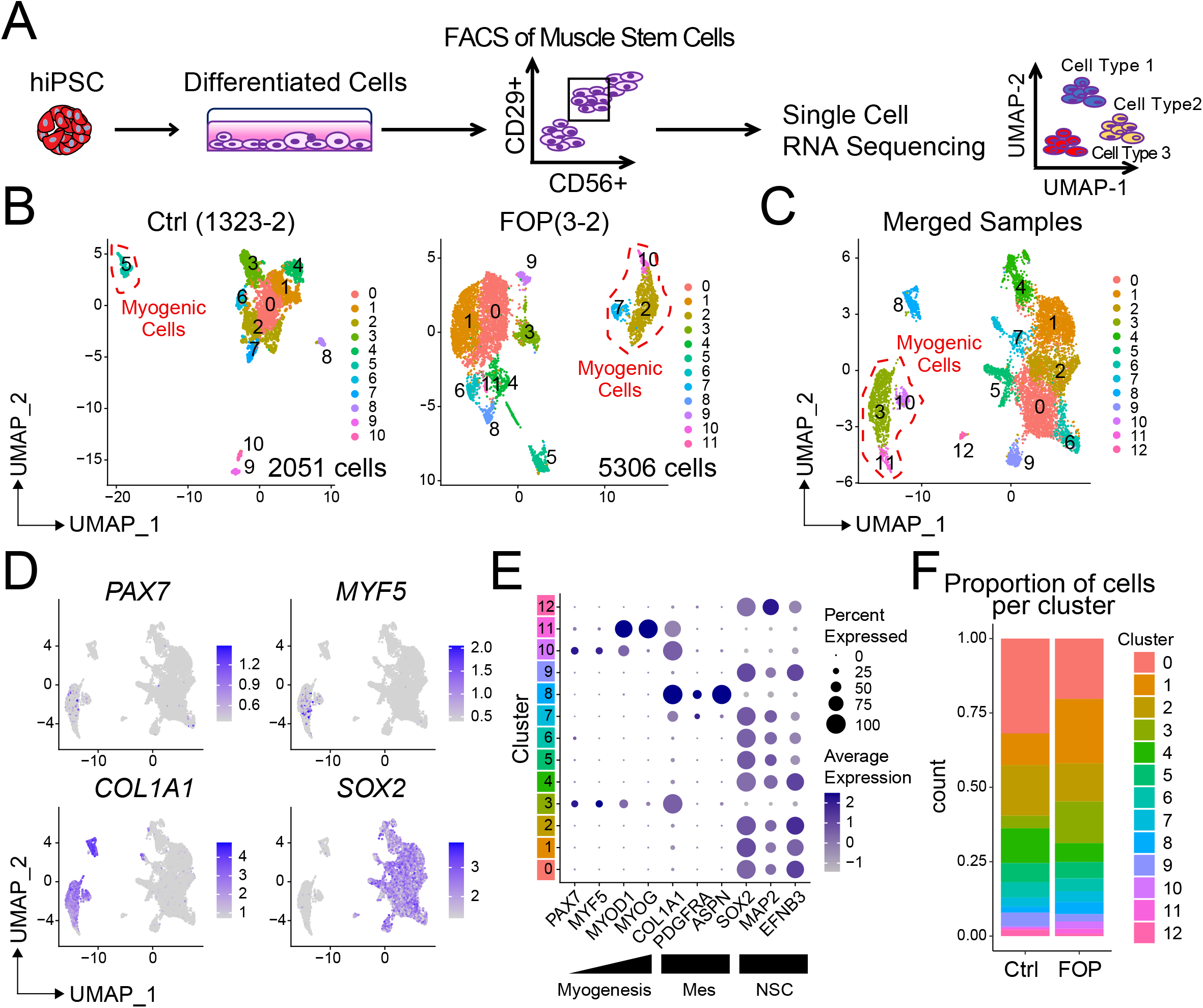
Single cell RNA sequencing of HNK1^-^ CD45-CD31^-^ CXCR4^+^CD29^+^ CD56^dim^ cells. **(A)** Schematic of differentiating hiPSCs, sorting of iMPCs, and scRNAseq. **(B)** UMAP visualization plots of the control (2,051 cells) and FOP (5,306 cells) samples. Each dot represents one cell colored by cluster identification. The myogenic cell type was assigned according to expression of a combination of marker genes **(Figure 4-figure supplement 1). (C)** UMAP of cells combined from both control and FOP samples. **(D)** Feature expression plots showing the localization of cells expressing myogenic markers, mesenchymal, and neural cell marker. (E) Dot plot displaying expression genes associated with myogenesis, neurogenesis and mesenchymal markers. **(F)** Proportion of cells per cluster for each sample.

### 6 FACS- sorted iMPC transcriptome is heterogeneous

Typical muscle progenitor cultures are expected to contain cells undergoing expansion, differentiation, and maturation. Sub-clustering (**Figure 5A,B**) identified 5 new myogenic subpopulations (**Figure 5B**). We ordered the myogenic cells into 3 major branches using the Monocle analysis package (Trapnell et al., 2014) and constructed pseudotime differentiation trajectories (**Figure 5C**). *PAX7 and MYF5* were upregulated in branch B but downregulated in branches A and C (**Figure 5C,D**). *MYOD* and *MYOG* showed higher expression patterns early in branch A and late in branch C (**Figure 5C,D**). Quiescent iMPCs were at the intersection of the 3 branches (clusters 0, 1) while distal parts of the 3 branches, notably comprised of clusters 3 and 4, contained more mature cells with transcriptional profiles similar to activated MuSCs, progenitor cells, or myoblasts. Cluster 4 expressed higher levels of *MYOD, MYOG, SOX8*, and *MEF2C* (markers of differentiated MuSCs/myoblasts/myocytes), while clusters 0-3 expressed higher levels of *PAX7* and *MYF5 (*markers of more quiescent MuSCs) (**Figure 5E,F**). Quiescence (*SPRY1, HEY1, H ES1*) and cell cycle (*TOP2A* or *KI67*) markers were increased in clusters 0-2 and cluster 3 respectively (**Figure 5F,G** and **Figure 5-figure supplement 1A,B**). Cluster 3 had a higher proportion of cells in G2M and S phase (**Figure 5-figure supplement 1B**). The cell cycle distribution was similar in control **Figure 5: Transcriptional profile of iMPCs. (A)** Identified myogenic clusters (in red) were sub-clustered from the other clusters (neurogenic and mesenchymal) and re-analyzed. **(B)** UMAP of the new myogenic subclusters was generated. **(C)** Pseudotime trajectory plot generated via Monocle analysis depicting all myogenic clusters. **(D)** Gene plots displaying the expression of specific myogenic genes as a function of pseudotime. Arrows represents the direction and major branches of the pseudotime. Branches A-C are marked with letters along the trajectories. **(E)** Feature expression of cells expressing myogenic markers. **(F)** Genes expression associated with stemness, quiescence, and activation for the described Hu-MuSCs subsets. **(G)** Feature expression plots of additional top differentially expressed marker for each cluster. **(H-I)** Violin plots of myogenic **(H)** and chondro/osteogenic genes **(I)** that were significantly differentially expressed. Violin plot width depicts the larger probability density of cells expressing each particular gene at the indicated expression level. *, significantly different following differential expression testing using the Wilcoxon rank sum test per cluster. **(J)** SMAD and P38MAPK pathway markers significantly differentially expressed between control and FOP myogenic cells. *p* values for **H-J** are in **Figure 5-Source Data 1**.

and FOP (**Figure 5-figure supplement 1C**). Cells expressing *APOE* and *KRT17* defined cluster 0. CDH15, a niche regulator of SCs quiescence (Goel, Rieder, Arnold, Radice, & Krauss, 2017) was enriched in cluster 1. Cluster 2 consisted of cells expressing high levels of *FOS* (**Figure 5F,G** and **Figure 5-figure supplement 1A**).

**Figure 5:**
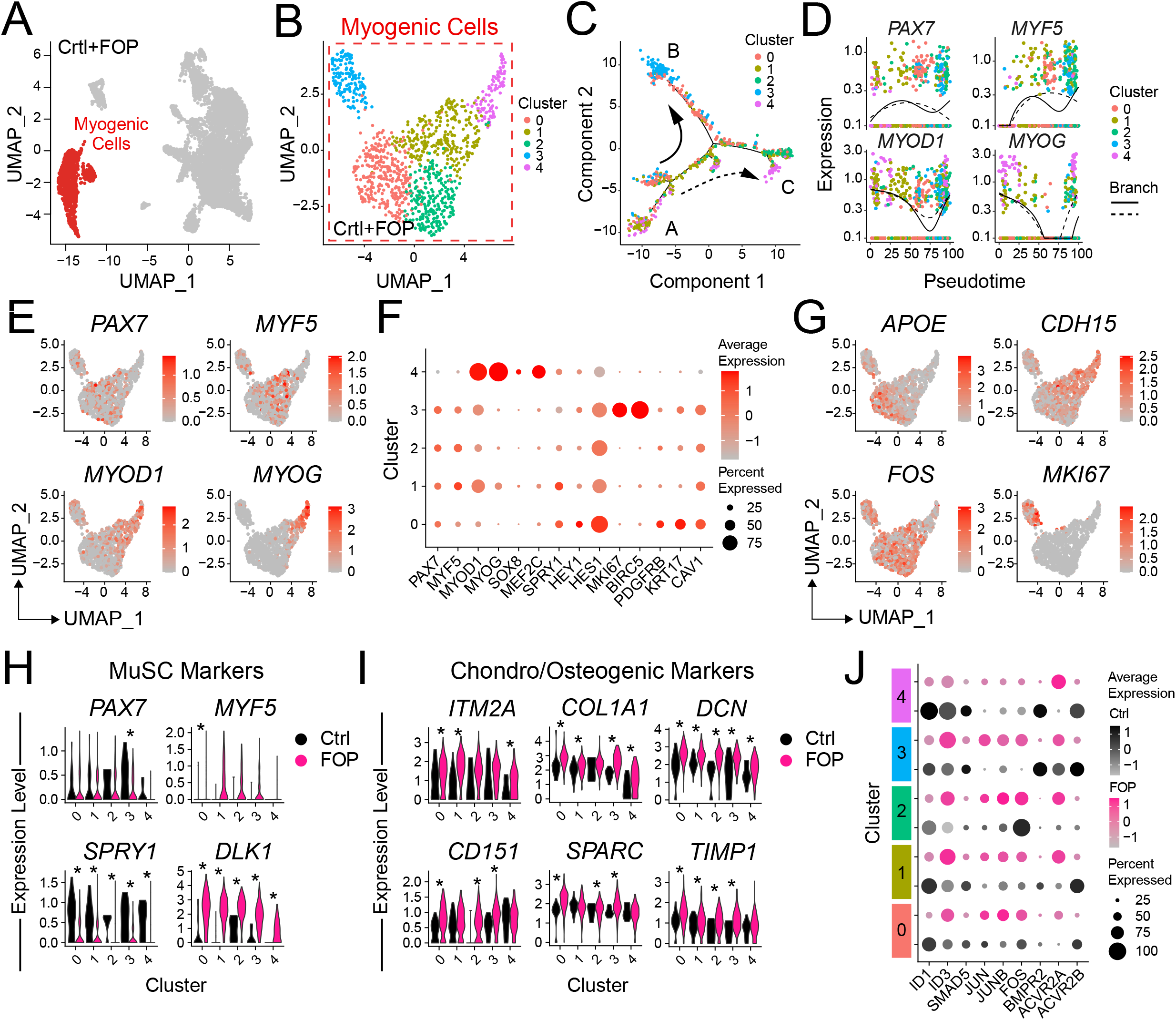
Transcriptional profile of iMPCs. **(A)** Identified myogenic clusters (in red) were sub-clustered from the other clusters (neurogenic and mesenchymal) and re-analyzed. **(B)** UMAP of the new myogenic subclusters was generated. **(C)** Pseudotime trajectory plot generated via Monocle analysis depicting all myogenic clusters. **(D)** Gene plots displaying the expression of specific myogenic genes as a function of pseudotime. Arrows represents the direction and major branches of the pseudotime. Branches A-C are marked with letters along the trajectories. **(E)** Feature expression of cells expressing myogenic markers. **(F)** Genes expression associated with stemness, quiescence, and activation for the described Hu-MuSCs subsets. **(G)** Feature expression plots of additional top differentially expressed marker for each cluster. **(H-I)** Violin plots of myogenic **(H)** and chondro/osteogenic genes **(I)** that were significantly differentially expressed. Violin plot width depicts the larger probability density of cells expressing each particular gene at the indicated expression level. *, significantly different following differential expression testing using the Wilcoxon rank sum test per cluster. **(J)** SMAD and P38MAPK pathway markers significantly differentially expressed between control and FOP myogenic cells. *p* values for **H-J** are in **Figure 5-Source Data 1**.

The proportion of cells in clusters 3 and 4 were similar in both control (10% and 12.7%) and FOP (14.3% and 9%) samples. The proportion of cells in clusters 0 and 3 was higher in FOP (31.4% and 23.4%) compared to control (20% and 5.5%), while the proportion of cells in cluster 2 was increased in control (51.8% vs 21.9%) (**Figure 5-figure supplement 1D**).

Thus, hiPSC differentiation cultures contain subpopulations of iMPCs showing the expected spectrum of quiescence, activation, and differentiation with FOP cultures having a higher proportion of cells in the stem cell/progenitor and proliferating phases and fewer mature myoblasts.

### 7 FOP iMPCs cells express increased chondro/osteogenic and ECM markers

Differential expression analysis on the transcriptional profiles of the sub-clustered myogenic cells (**Figure 5A,B** and **Figure 5-figure supplement 1E**) was used to see if ACVR1^R206H^ altered transcriptional signatures. While *PAX7* was significantly increased in control vs. FOP cells in cluster 3 only (**Figure 5H**), *MYF5* was significantly increased in FOP cells from cluster 1 compared to control cells (**Figure 5-Source Data 1**). Since Hu-MuSCs show heterogeneous levels of PAX7 and MYF5 expression (Kuang et al., 2007), this suggests the ACVR1^R206H^ mutation may favor one sub-population over another. Interestingly, *SPRY1*, a known regulator of quiescence (Shea et al., 2010) which decreases with age (Bigot et al., 2015), was significantly downregulated in FOP cells of clusters 0-4, while *DLK1*, which act as a muscle regeneration inhibitor (Andersen et al., 2013) was significantly increased in FOP cells (**Figure 5H** and **Figure 5-Source Data 1**).

Since the ACVR1^R206H^ mutation increases BMP pathway activity and expression of chondrogenic and osteogenic markers in multiple lineages (Barruet et al., 2016; Culbert et al., 2014; Matsumoto et al., 2013), we examined the expression levels of extracellular matrix, fibrogenic, chondrogenic, and osteogenic genes. *ITM2A, COL1A1, DCN, CD15, SPARC*, and *TIMP1* were significantly increased in FOP cells (**Figure 5I** and **Figure 1-Source Data 1**). ECM proteoglycans known to be involved in inflammation, including *BGN* (Nastase, Young, & Schaefer, 2012) and *LUM* (Nikitovic, Papoutsidakis, Karamanos, & Tzanakakis, 2014), *TAGLN* [regulates osteogenic differentiation (Elsafadi et al., 2016)], and *IGBP5* [increased in aged satellite cells (Soriano-Arroquia, McCormick, Molloy, McArdle, & Goljanek-Whysall, 2016)], were also increased in FOP cells (**Figure 5-figure supplement 1F** and **Figure 1-Source Data 1**).

Finally, gene expression of target genes of the BMP pathway (*ID1, ID3, BMPs*, and *SMADs*) and the p38MAPK pathway (**Figure 5J** and **Figure 5-figure supplement 1G**) was assessed to see if activated ACVR1 altered these pathways. *ID1* was significantly higher (clusters 0, 1, 3, and 4) in control cells while *ID3* was significantly higher in FOP cells (clusters 1, 2). In addition, the BMP/TGFβ pathway downstream target gene *SMAD5* was significantly higher in cluster 2 of control cells. The p38 pathway components *JUN* (clusters 1, 3), *JUNB* (clusters 0-2), and *FOS* (clusters 1, 3) were significantly increased in FOP cells. Within the known ACVR1 co-receptors, *BMPR2* (clusters 3, 4) and *ACVR2B* (clusters 0, 2, 3) expression were significantly higher in control cells while *ACVR2A* expression was significantly higher in FOP cells (cluster 1, 4). *ACVR2B* was significantly higher in control cells (cluster 0, 1, 4) (**Figure 5J**). Similar to the primary FOP Hu-MuSCs (**Figure 1**), these results suggest that FOP iMPCs have a chondrogenic/osteogenic signature, increased *ID3* expression, and also showed higher p38 pathway activity and higher levels of the *ACVR2A* co-receptor at different stages of myogenic differentiation.

### 8 iMPC transcriptome shows similarities to primary Hu-MuSCs

Comparing the iMPC scRNAseq to primary Hu-MuSCs data of sorted satellite cells from a human vastus lateralis muscle (Barruet et al., 2020) was used to identify if their lower engraftment efficiency was due to transcriptional differences. The merged data UMAP (**Figure 6A**) showed that clusters 0, 1, 4, 6, and 7 contained myogenic cells (**Figure 6B,C**). *PAX7*^*+*^ cells were identified in clusters 0, 1, 4, and 7 (**Figure 6C**). Myocyte contaminants present in the primary sorted cells constituted cluster 6. Although iMPCs expressed *PAX7* and *MYF5*, the gene expression levels were higher in primary Hu-MuSCs. In contrast, *MYOD* was higher in iMPCs (**Figure 6C** and **Figure 6-figure supplement 1**).

**Figure 6:**
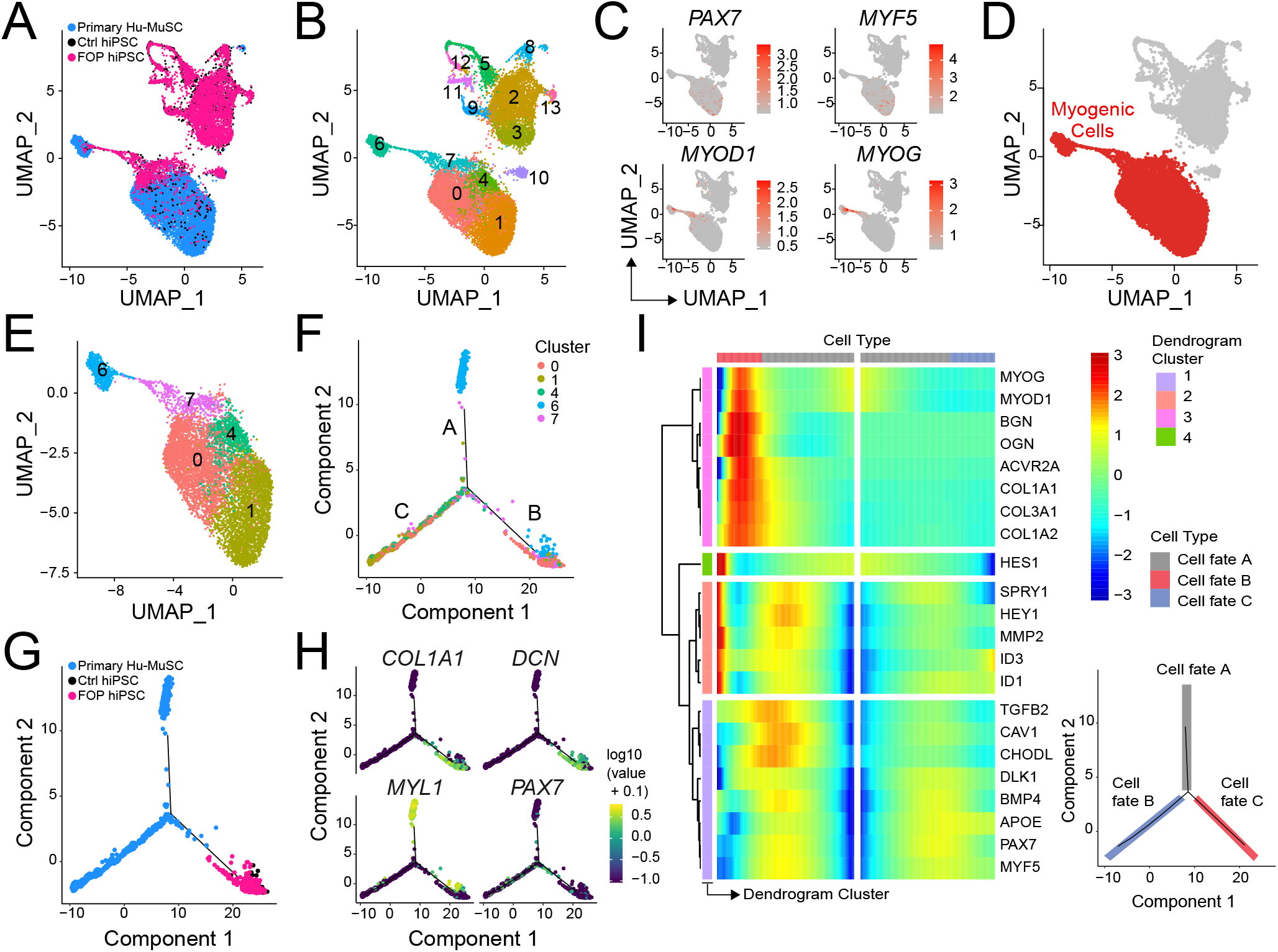
iMPC transcriptional signature compared to human primary muscle stem cells. **(A-B)** UMAP of cells combined from human primary muscle stem cells (vastus muscle) and the control and FOP samples. **(A)** UMAP showing the distribution of cells per sample, and **(B)** with clusters labeled. **(C)** Feature expression plots of cells expressing myogenic markers. **(D)** Myogenic cells (in red) are comprised in cluster 0, 1, 4, 6, and 7. **(E)** UMAP of the myogenic clusters only used for the pseudotime analysis. **(F-G)** Pseudotime trajectory plot generated via Monocle analysis depicting all myogenic clusters **(F)** and samples **(G)**. Branches are annotated A-C in **(F). (H)** Level of expression of ECM/osteogenic genes along the cell trajectories. **(I)** Heatmap representing genes that are significantly branch dependent using the BEAM analysis (**Figure 6-Source Data 1**) and also genes that have similar lineage-dependent expression patterns. Cell fate (branches) are shown in the lower right panel.

**Figure 7.**
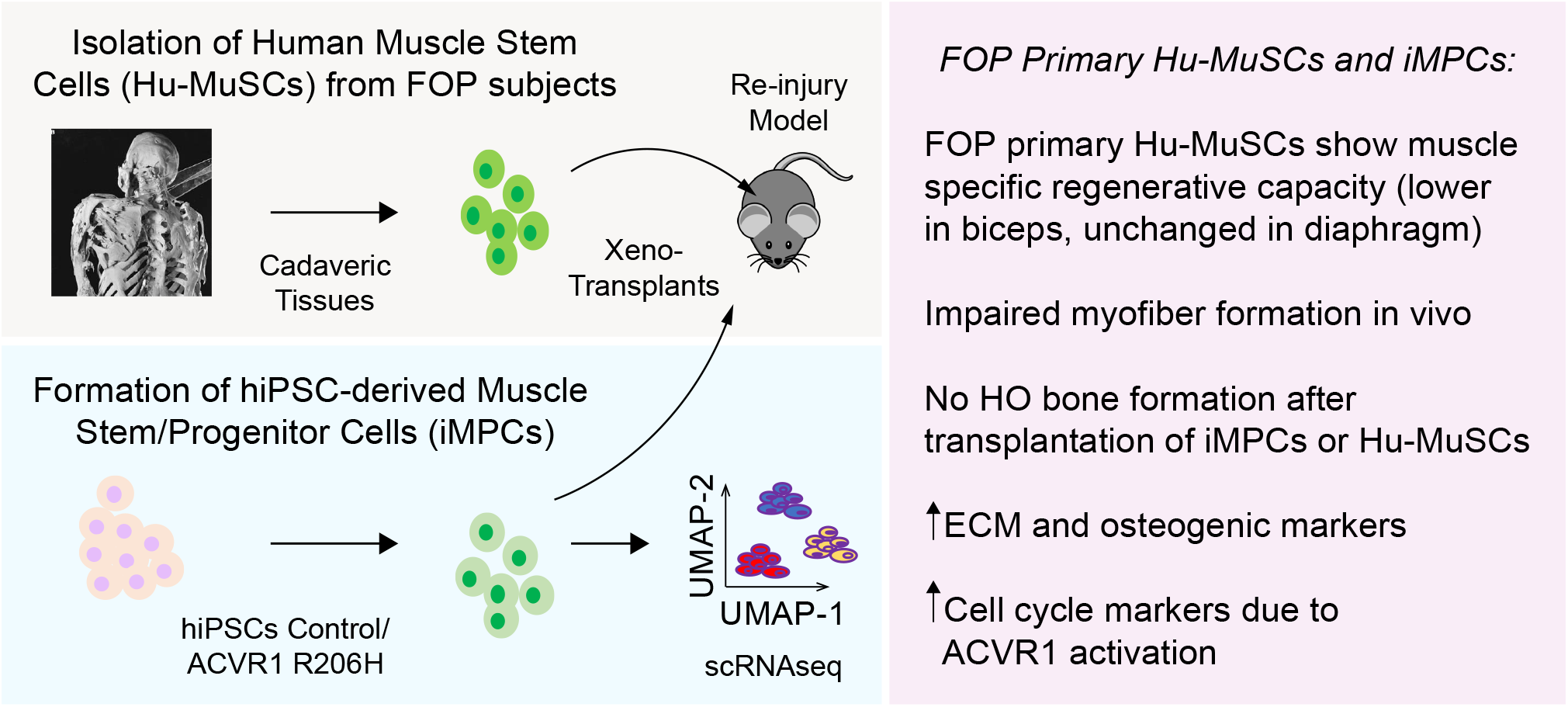
Testing the role of activated ACVR1 signaling in muscle repair. FOP iMPCs and FOP primary human muscle stem cells are used for modeling muscle stem cell engraftment and regeneration properties.

Detailed analysis of the myogenic cell subset (cluster 0, 1, 4, 6 and 7, **Figure 6D**, cells labeled in red, and **Figure 6E**) was done using pseudotime trajectory analysis to elucidate the states of the iMPCs with respect to primary Hu-MuSCs. Cells from clusters 0, 1, and 4 were distributed along branches B and C. Myocytes ordered at the distal end of branch A. iMPCs ordered away from primary Hu-MuSCs (*PAX7*+) and primary myocytes (*MYL1*+) (**Figure 6F-H**). Branch expression analysis modeling (BEAM) allowed us to investigate significant gene that are branch-dependent in their expression (**Figure 6-Source Data 1**). Branch B, consisting of iMPCs and subset of primary Hu-MuSCs, expressed significantly higher levels of genes associated with mesenchymal, fibrogenic, chondrogenic, osteogenic lineages and extracellular matrix (**Figure 6H,I**). Thus, iMPCs retained strong bi-potency compared to primary Hu-MuSCs, suggesting that iMPCs may not be as committed to the muscle lineage as primary adult muscle stem cells (Xi et al., 2020).

## Discussion

Developing optimal strategies for skeletal muscle regeneration and repair requires a detailed understanding of how these processes are regulated. Especially for diseases like FOP where muscle injury can trigger disease progression or severe complications, generating large numbers of human iMPCs using transgene-free protocols, holds promise for understanding pathological mechanisms and finding new therapeutic targets. This study revealed several novel findings, including deficiencies in the muscle repair capacity of FOP Hu-MuSCs, abnormal production of extracellular matrix components by iMSCs that likely contribute to FOP disease progression, and important differences between primary Hu-MuSCs and iMPCs that may impact other studies. In addition, our results suggest that there are muscle specific differences in Hu-MuSC repair capacity, potentially explaining why some skeletal muscles appear to be protected from developing HO.

The iMPC and primary Hu-MuSC single cell transcriptomes revealed clear subsets of cells at all stages of muscle differentiation (quiescent, activated, and differentiated). Satellite cell subtype markers [e.g. *COLs, DLK1, ID3*, and *HES1*, (Barruet et al., 2020)] were also highly expressed in iMPCs suggesting that *in vitro* differentiation may favor specific subtypes of satellite cells. iMPCs also expressed high levels of mesenchymal/ECM markers, similar to that in human fetal muscle progenitor cells (Xi et al., 2020). These results suggest that iMPCs may retain a more progenitor-like phenotype as compared to Hu-MuSCs. In addition, we showed that PAX7^+^ iMPCs can engraft in muscle injury models in mice, despite heterogeneity in the cultures and lower expression of typical satellite cells markers (e.g. *PAX7, MYF5*).

This iMPC system revealed that activation of the BMP pathway by the FOP ACVR1^R206H^ mutation induced changes that could contribute to abnormal muscle healing. As expected, the classical SMAD pathways were active in FOP cells; however, it was unexpected to find that the p38 pathway via JUN/FOS was more active in the FOP iMPCs, considering that FOP cultures had a higher proportion of cells in the stem cell/progenitor and proliferating phases. While increased SMAD activity (Billings et al., 2008) or misinterpretation of the Activin A ligand (Hatsell et al., 2015) occur with ACVR1^R206H^ mutation, abnormal p38 signaling may be critical in some cells like macrophages (Barruet et al., 2018). Since p38 is a major regulator of Hu-MuSC function (Segales, Perdiguero, & Munoz-Canoves, 2016) and has been associated with inflammation and ECM accumulation in aged muscle (Cosgrove et al., 2014), further studies are needed to determine how this p38 signaling contributes to satellite cell subtype specification (Barruet et al., 2020) and affect healing in FOP.

Importantly, our iMPC and Hu-MuSC transplant studies showed no HO in the recipient mice, consistent with prior FOP genetic studies (Dey et al., 2016; Lees-Shepard et al., 2018). However, FOP iMPCs showed increased chondrogenic/osteogenic and ECM gene expression, a feature seen in other bone-related cell types in FOP (Barruet et al., 2016; Culbert et al., 2014; Lees-Shepard et al., 2018). Thus, FOP Hu-MuSCs may contribute to HO formation indirectly by modulation of the osteogenic environment such as contributing to changes in muscle stiffness, as previously reported in mice (Stanley, Heo, Mauck, Mourkioti, & Shore, 2019). In addition, while our results suggest no major differences or possibly a slight increase in the ability to form skeletal muscle progenitors, activated ACVR1 by the R206H mutation decreased (but did not abrogate) in vitro formation of mature myoblasts and in vivo muscle repair after transplant. This impaired formation of mature skeletal muscle likely contributes to the abnormal skeletal muscle healing in FOP and may predispose patients with FOP to HO formation.

Comparing primary FOP Hu-MuSCs from muscles that develop HO (biceps) and non-affected muscle (diaphragm) suggests that the source of the Hu-MuSCs impacts engraftment efficiency. The re-injury model showed that engraftment of Hu-MuSCs from FOP biceps, but not from diaphragm, remained significantly decreased. Thus, it is intriguing to consider that the clinical sparing of the diaphragm from HO in patients with FOP may result from a less impaired or unimpaired muscle repair process in diaphragm satellite cells. Further delineation of muscle-specific Hu-MuSC properties will be revealing to understand this observation. Our finding that primary FOP Hu-MuSCs have lower engraftment ability provides a potential explanation for the poor skeletal muscle repair observed in patients with FOP (Shore, 2012). However, the FOP and control iMPCs showed no major differences in engraftment, possibly due to decreased assay sensitivity from the lower engraftment efficiency of iMPCs in general, because iMPCs may be more immature than primary Hu-MuSCs, or that iMPCs represent a subtype of cells that may be more reflective of non-ossifying skeletal muscle like diaphragm.

This study has several limitations. Our transplant experiments used immunocompromised mice where mature B and T cells are absent and macrophages are defective (Shultz et al., 2005). This may dampen the engraftment/regenerative phenotypes, particularly as immune cells are important in muscle regeneration (Furrer & Handschin, 2017) and there is growing awareness of the role of the immune system in the pathogenesis of FOP (Barruet et al., 2018; Convente et al., 2018). Also, while our FACS strategy can isolate Hu-MuSCs and iMPCs without the need of transcription factors or markers, we noted that the engraftment efficiency still varied among the iMPC lines despite FACS purification. This heterogeneity is an ongoing problem among all published iMPC protocols to date. In our case, this was counter-balanced by our consistent findings across multiple lines and between iMPCs and the availability of primary Hu-MuSCs. Although our primary cell studies were limited by the rarity of FOP (estimated at 1 in 1.4 million people) and even rarer suitable cadaveric samples, the combination of iPSC-derived lineages with the rare primary samples provided multiple avenues for supporting our conclusions. Finally, the ACVR1^R206H^ mutation increases BMP signaling but also introduces neofunction to Activin A. Our studies are not able to distinguish between these two contributing pathways. Our finding of increased p38 activity in the FOP iMPCs was also seen in FOP subject monocyte-derived macrophages (Barruet et al., 2018), suggesting that this alternate signaling pathway by ACVR1 should also be investigated further. Future studies will help elucidate the different factors that contribute to the hiPSC-line specific effects, including potential roles for disease modifier genes or BMP pathway modulators.

This study shows that human iPSC-derived muscle stem cells can be a valuable tool to model musculoskeletal diseases of skeletal muscle injury and repair. Correlating the findings in iMPCs with primary Hu-MuSCs revealed an indirect role for skeletal muscle progenitors in HO formation, as well as subtypes of Hu-MuSCs that may contribute to skeletal muscle specific regenerative capacity. These studies highlight the importance of skeletal muscle regeneration in disease pathogenesis and establish a foundation for understanding how skeletal muscle repair and osteogenesis are linked.

## Materials and Methods

| **Key Resources Table** |  |  |  |  |
| --- | --- | --- | --- | --- |
| **Reagent type (species) or resource** | **Designation** | **Source or reference** | **Identifiers** | **Additional information** |
| NSG mice | NOD.Cg-Prkdcscid Il2rgtm1Wjl/SzJ | https://www.jax.org/strain/005557 | 005557 | 8-12 week-old |
| Sequenced-based reagent | Human RT-PCR Primers | Applied Biosystems Taqman Assays | B-ACTIN<br>Hs01060665_g1 |  |
| Sequenced-based reagent | Human RT-PCR Primers | Applied Biosystems Taqman Assays | DYSTROPHIN<br>Hs007758098_m1 |  |
| Sequenced-based reagent | Human RT-PCR Primers | Applied Biosystems Taqman Assays | PAX7<br>Hs00242962_m1 |  |
| Sequenced-based reagent | Human RT-PCR Primers | Applied Biosystems Taqman Assays | CD56<br>HS00941830_m1 |  |
| Sequenced-based reagent | Human RT-PCR Primers | Applied Biosystems Taqman Assays | MYOGENIN<br>Hs01072232_m1 |  |
| Sequenced-based reagent | Human RT-PCR Primers | Applied Biosystems Taqman Assays | COL1A1<br>Hs01076780_g1 |  |
| Sequenced-based reagent | Human RT-PCR Primers | Applied Biosystems Taqman Assays | ID1<br>Hs03676575_s1 |  |
| Sequenced-based reagent | Human RT-PCR Primers | Applied Biosystems Taqman Assays | ID3<br>Hs00954037_g1 |  |
| antibody | Mouse monoclonal anti-Human DYSTROPHIN | DSHB | MANDYS104(7F7) | IF(1:10) |
| antibody | Mouse monoclonal anti-Human/Mouse DYSTROPHIN | Thermofisher | PA5-16734 | IF(1:500) |
| antibody | Mouse monoclonal anti-Human/Mouse PAX7 | DSHB | PAX7 | IF(1:10) |
| antibody | Mouse monoclonal anti-Human/Mouse MHC | DSHB | MF20 | IF(1:100) |
| antibody | Rabbit polyclonal anti-LAMININ | Sigma-Aldrich | L9393 | IF(1:250) |
| antibody | Mouse monoclonal anti-Human SPECTRIN | Leica Microsystems | NCL-SPEC1 | IF(1:100) |
| antibody | Mouse monoclonal anti-Human LAMIN A/C | Vector Laboratories | VP-L550 | IF(1:100) |
| antibody | Mouse monoclonal anti-Human CD31 (Beads) | Miltenyi Biotec | 130-091-935 | FACS |
| antibody | Mouse monoclonal anti-Human CD45 (Beads) | Miltenyi Biotec | 130-045-801 | FACS |
| antibody | Mouse monoclonal anti-Human CD31 AF450 (WM-59) | Ebioscience | 48-0319-42 | FACS |
| antibody | Mouse monoclonal anti-Human CD34 eFluor450 (4H11) | Ebioscience | 48-0349-42 | FACS |
| antibody | Mouse monoclonal anti-Human CD45 AF450 (30-F11) | Ebioscience | 48-0451-82 | FACS |
| antibody | Mouse monoclonal anti-Human CD29 FITC (TS2/16) | Ebioscience | 11-0299-41 | FACS |
| antibody | Recombinant human anti-CD56 APC-vio-770 (REA196) | Miltenyi Biotec | 130-114-548 | FACS |
| antibody | Mouse monoclonal anti-Human CXCR4 PE (12G5) | Ebioscience | 12-9999-41 | FACS |
| antibody | Mouse monoclonal anti-Human HNK1 PE (TB01) | Ebioscience | 12-057742 | FACS |
| antibody | Mouse monoclonal anti-Human CD45 PE (30-F11) | Ebioscience | 12-04551-82 | FACS |
| antibody | Mouse monoclonal anti-Human CD31 PE (390) | Ebioscience | 12-0311-82 | FACS |
| antibody | Mouse monoclonal anti-Human CD56 APC (CMSSB) | Ebioscience | 47-0567-42 | FACS |
| antibody | Mouse monoclonal anti-Human CXCR4 PE-Cy7 (12G5) | Ebioscience | 25-9999-42 | FACS |
| antibody | FcR block | Miltenyi Biotec | 130-059-901 |  |
| software, algorithm | GraphPad Prism | GraphPad Prism (https://graphpad.com) |  |  |
| software, algorithm | Seurat (3.1.5) | https://satijalab.org/seurat/ |  |  |
| software, algorithm | Monocle (2.12.0) | http://cole-trapnell-lab.github.io/monocle-release/ |  |  |
| software, algorithm | cellranger | https://support.10xgenomics.com/single-cell-gene-expression/software/pipelines/latest/feature-bc |  |  |
| software, algorithm | FlowJo | https://www.flowjo.com |  |  |

### Cell Culture and Differentiation

Pluripotent hiPSC lines derived from control and FOP fibroblasts (Matsumoto et al., 2013; Spencer et al., 2014) were cultured in mTeSR1 medium (StemCell Technologies) on irradiated SNL feeder cells (McMahon & Bradley, 1990) as described previously. hiPSCs were passaged at least once on Matrigel (Corning)-coated plates (150-300µg/ml) to remove the SNLs before use in differentiation assays. ROCK inhibitor Y-27632 (10µM, StemCell Technologies) was added to mTeSR1 when cells were split and removed the following day.

hiPSC lines were differentiated into skeletal muscle cells using modifications based on prior protocols (Chal et al., 2016; Shelton et al., 2014). Our hiPSC lines differentiated better with a lower cell number seeding and a longer time of recovery between the seeding and the start of the differentiation (2 days, data not shown). Cells were seeded at 7.5×10^5^ cells per well of a 12-well plate on Matrigel two days before the differentiation medium (E6 medium, supplemented with either 10µM CHIR99021 (Tocris) for 2 days or with 3µM CHIR99021 and 0.5µM LDN193189 for six days). Cells were then grown in un-supplemented E6 media until day 12, then changed to StemPro-34 media supplemented with 10ng/ml bFGF until day 20. The medium was then replaced by E6 medium until day 35, when DMEM/F12 supplemented with N2 (Gibco) and Insulin-Transferrin-Selenium (ITS-A, 100X Gibco) was added. Media was changed daily until harvest at day 50. After sorting, cells were plated in satellite cell media [DMEM/F12 (Gibco), 20% FBS (Hyclone), 1X ITS (Gibco), 1X Penicillin/ Streptomycin (Gibco)] for further functional assays (**Figure 2A,F**). Once cells reached confluence, cells were cultured in differentiation media (DMEM, 2% horse serum, 1X Penicillin/ Streptomycin; Gibco).

### Flow cytometry

HNK1^-^CD45^-^CD31^-^CXCR4^+^CD29^+^CD56^dim^ cells were sorted from skeletal muscle differentiation of control and FOP hiPSCs (described in detail in Supplemental Experimental Procedures). Human primary satellite cells were isolated and sorted as described (Garcia et al., 2018; Garcia et al., 2017). Human muscle was freshly harvested and stored in DMEM with 30% FBS at 4°C overnight or for 2 extra days (delay due to shipping). Muscle samples were digested, erythrocytes were lysed, and hematopoietic and endothelial cells were depleted with magnetic column depletion using CD31, CD34, and CD45 (eBioscience). Cells were further gated as described in **Figure 1-figure supplement 1A** and sorted for CXCR4^+^/CD29^+^/CD56^+^ and collected for subsequent experimentation.

### Animal care and Transplantation studies

All mouse studies were performed using protocols approved by the UCSF Institutional Animal Care and Use Committee. Mice were either bred and housed in a pathogen-free facility at UCSF or purchased from The Jackson Laboratory. 8-12 week-old NSG (NOD.Cg-Prkdcscid Il2rgtm1Wjl/SzJ) mice were randomized to all experimental groups by sex and littermates. The TA of each mice was irradiated with 18 Gy before transplantation. Isolated primary human satellite cells (Hu-MuSCs) or iMPCs were injected with 50 µl 0.5% bupivacaine directly into the TA muscle of one leg as described (Garcia et al., 2018; Garcia et al., 2017) and summarized in the Supplemental Experimental Procedures. The TA for each mouse was harvested at week 5 or week 10 after transplantation and frozen in O.C.T. compound in 2-methylbutane chilled in liquid nitrogen. Serial 6µm transverse frozen sections were analyzed or stored at −80°C.

### Cell Immunostaining and NSG Tibialis Anterior Analysis

iMPCs were fixed with 4%PFA/PBS for 10min at room temperature, permeabilized with 0.1%Triton-100X (Sigma-Aldrich), and blocked with 5% BSA (Sigma-Aldrich). Cells were stained overnight with primary antibodies for PAX7, MYOGENIN, DYSTROPHIN, and MHC. Cells were then incubated for 1 hr at room temperature in the dark with secondary antibodies Alexa488-conjugated goat anti-mouse IgG and Alexa546-conjugated goat anti-mouse IgG (Invitrogen). Nuclei were stained with DAPI (Sigma-Aldrich). Images were taken on a Nikon Eclipse E800 or Leica DMI 4000B.

### Immunohistochemistry and Immunofluorescence of Human Muscle Samples

Human muscle samples were fixed in neutral buffered formalin for 24 h and then placed in 70% ethanol for at least 24 h. The sample with heterotopic bone was decalcified in 10% EDTA (pH 7.2-7.4) before paraffin embedding and sectioning. Sections were stained with hematoxylin and eosin (J. David Gladstone Institutes Histology Core) or for alcian blue (pH 1.0) for cartilage and nuclear red stain for nuclei.

Freshly harvested human muscle was stored in DMEM with 30% FBS at 4°C, or snap in frozen in O.C.T. compound in 2-methylbutane chilled in liquid nitrogen. Serial 6µm transverse frozen sections were analyzed or stored at −80°C and processed similarly to the mouse TA samples above. Sections were stained with PAX7 (DSHB) and mouse monoclonal anti-Collagen Type I (Millipore-Sigma). Details about specimens are in **Figure 1-Data Source 1**.

### RT-PCR and Quantitative Analysis

Tissues were collected in TRI Reagent (Sigma-Aldrich) to isolate total RNA using the Arcturus™ PicoPure™ RNA kit (Applied Biosystems) as previously described for small samples (Schepers, Hsiao, Garg, Scott, & Passegue, 2012). 0.2 to 0.5 µg of RNA were transcribed into cDNA with VeriScript cDNA synthesis kit (Affymetrix). cDNA was then pre-amplified with GE PreAmp Master Mix (Fluidigm Inc). Real-time quantitative PCR was performed in triplicated with either VeriQuest Probe qPCR Master Mix (Affymetrix) or Taqman Universal PCR Master Mix (Life Technologies) on either a Viia7 thermocycler (Life Technologies) or on a BioMark 48.48 dynamic array nanofluidic chip (Fluidigm, Inc) according to manufacturers’ instructions. Beta actin was used for normalization as endogenous control.

### Single cell RNA Sequencing and Analysis

scRNAseq was performed using the Chromium Single Cell 3’ Reagent Version 2 Kit from 10X Genomics. 45,000 (FOP) and 30,000 (control) HNK1^-^CD45^-^CD31^-^ CXCR4^+^CD29^+^CD56^dim^ cells isolated from iMPC differentiations were analyzed, using the sequencing procedures and analytical strategies described in Supplemental Experimental Procedures.

### Statistical analysis

The data were analyzed utilizing GraphPad Prism v.7 software (GraphPad) using one-way (transplant) and two-way ANOVA with post hoc Tukey’s or Sidak’s multiple comparison test (gene expression). The Sidak test was used when comparing means between control and FOP, and the Tukey test was used when means of both control and FOP were compared together with other groups for the gene expression data. For the transplantation studies, at least 3 mice were used per group. At least three biological replicates were performed for each experiment unless indicated otherwise. All error bars are depicted as standard deviation, p-values are (*p < 0.05, **p < 0.01, ***p < 0.001, ****p < 0.0001).

### Human Specimen Procurement

Human samples were collected through the UCSF Biospecimens and Skeletal Tissues for Rare and Orphan Disease Genetics (BSTROnG) Biobank, using protocols approved by the UCSF Institutional Review Board. All participants provided written consent. For the transplantation studies (**Figure 6D-H** and **Figure 6-figure supplement 1**), biopsy of the control subject was obtained from a 44 yo female healthy individual undergoing surgery at UCSF, the muscle from the FOP patient was obtained at autopsy from a 55 yo. Written informed consent was obtain from all subjects or their families.

## Acknowledgments

The authors thank Kelly Wentworth and Samuel Kou for their assistance collecting the autopsy samples and Francesco Tedesco for his support on the AFM grant. This work was supported by a NIH/NIAMS R01AR066735 to ECH, a French Muscular Association (AFM-Telethon) Trampoline grant to ECH and EB, the Radiant Hope Foundation to ECH, and the UCSF Cohort Development Grant to ECH; the California Institute for Regenerative Medicine Fellowship Program to UCSF (TG2-01153) to EB, and the UCSF Program for Breakthrough Biomedical Research (PBBR) to EB; and the NIH R01AR072638-03 to JHP. Finally, the authors would like to thank the patients and their families for their generous specimen donations.

## Author contributions

Conceptualization, ECH, JHP, EB; Methodology, EB; Software Analyses, EB, TM; Validation, EB, SMG; Formal Analysis, EB; Investigation, EB, SMG, JW, BMM, ST; Resources, ECH, JHP; Data Curation, EB, TM; Writing – Original Draft, EB, ECH; Writing – Review and Editing, EB, ECH, JHP; Visualization, EB; Supervision, ECH, JHP; Project Administration, ECH, EB; Funding, ECH, EB, JHP; clinical samples, ECH, JHP.

## Data and materials availability

Single cell gene expression data have been deposited (GSE151918). The dataset used for the primary Hu-MuSCs can be found here, https://datadryad.org/stash/landing/show?id=doi%3A10.7272%2FQ65X273X. Detailed scripts can be found here, https://github.com/EmilieB12/FOP_muscle/tree/main.

## Abbreviations

PAX7: Paired Box 7
hiPSCs: Human induced pluripotent stem cells
BMP: bone morphogenetic protein
ACVR1/2A: activin receptor type
1/2A MYOD1: Myogenic Differentiation 1
HNK1: Human Natural Killer-1
CXCR4: C-X-C chemokine receptor type 4
MYF5: Myogenic Factor 5
COL1A1: Collagen Type I Alpha 1 Chain
PDGFRA: Platelet Derived Growth Factor Receptor Alpha
SOX2: SRY-Box 2
EFNB3: Ephrin B3
MAP2: Microtubule Associated Protein 2
ASCL1: Achaete-Scute Family BHLH Transcription Factor 1
ASPN: Asporin
APOE: Apolipoprotein E
OGN: Osteoglycin
TOP2A: DNA Topoisomerase II Alpha
FABP7: Fatty Acid Binding Protein 7
SPRY1: Sprouty 1
MYL1: Myosin Light Chain 1
DLK1: Delta Like Non-Canonical Notch Ligand 1
ITM2A: Integral Membrane Protein 2A
DCN: Decorin
SPARC: Secreted Protein Acidic and Cysteine Rich
TIMP1: Tissue Inhibitor of Metalloproteinase 1
BGN: Biglycan
LUM: Lumican
TAGLN: Transgelin
IGBP5: Insulin Like Growth Factor Binding 5
ID1/3: Inhibitor of DNA Binding 1/3
TGFb: Transforming Growth Factor b
BMPR2: Bone Morphogenetic Protein Receptor 2
CAV1: Caveolin 1
ECM: Extracellular Matrix

## Supplementary Materials

**Video 1. Myotube contraction of differentiated Control (1323-2) hiPSCs**.

**Video 2. Myotube contraction of differentiated FOP (F1-1) hiPSCs**.

**Data File S1- hiPSCs samples integration**.

**Data File S2- Myogenic sub-clustering**.

**Data File S3- Pseudotime analyses**.

**Data File S4- Primary Hu-MuSCs/hiPSCs integration**.

**Figure 1-figure supplement 1.**
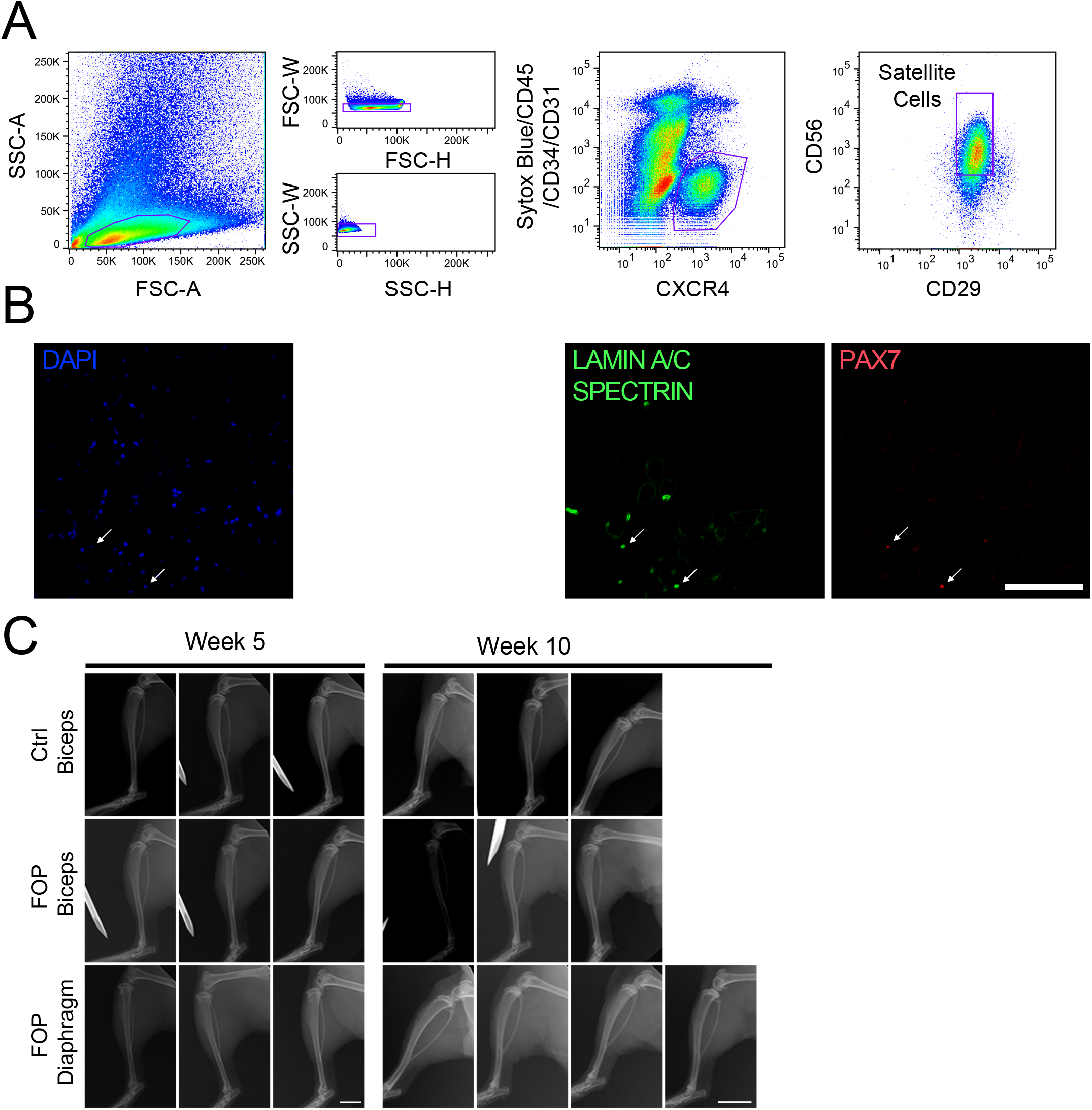
Transplanted satellite cells from biceps and diaphragm of FOP patient do not form bone. **(A)** Flow cytometry analysis sorting strategy to isolate human satellite cells. **(B)** Representative immunostaining of human satellite cells. White arrows indicate PAX7+ satellite cells (scale bar, 50µm). **(C)** X-ray of transplanted mice at 5 and 10 weeks after initial transplantation (scale bars, 4mm (week 5), 7mm (week 10)) showing no heterotopic ossification. Details about muscle specimens used in this figure are in **Figure 2-Source Data1**.

**Figure 2-figure supplement 1.**
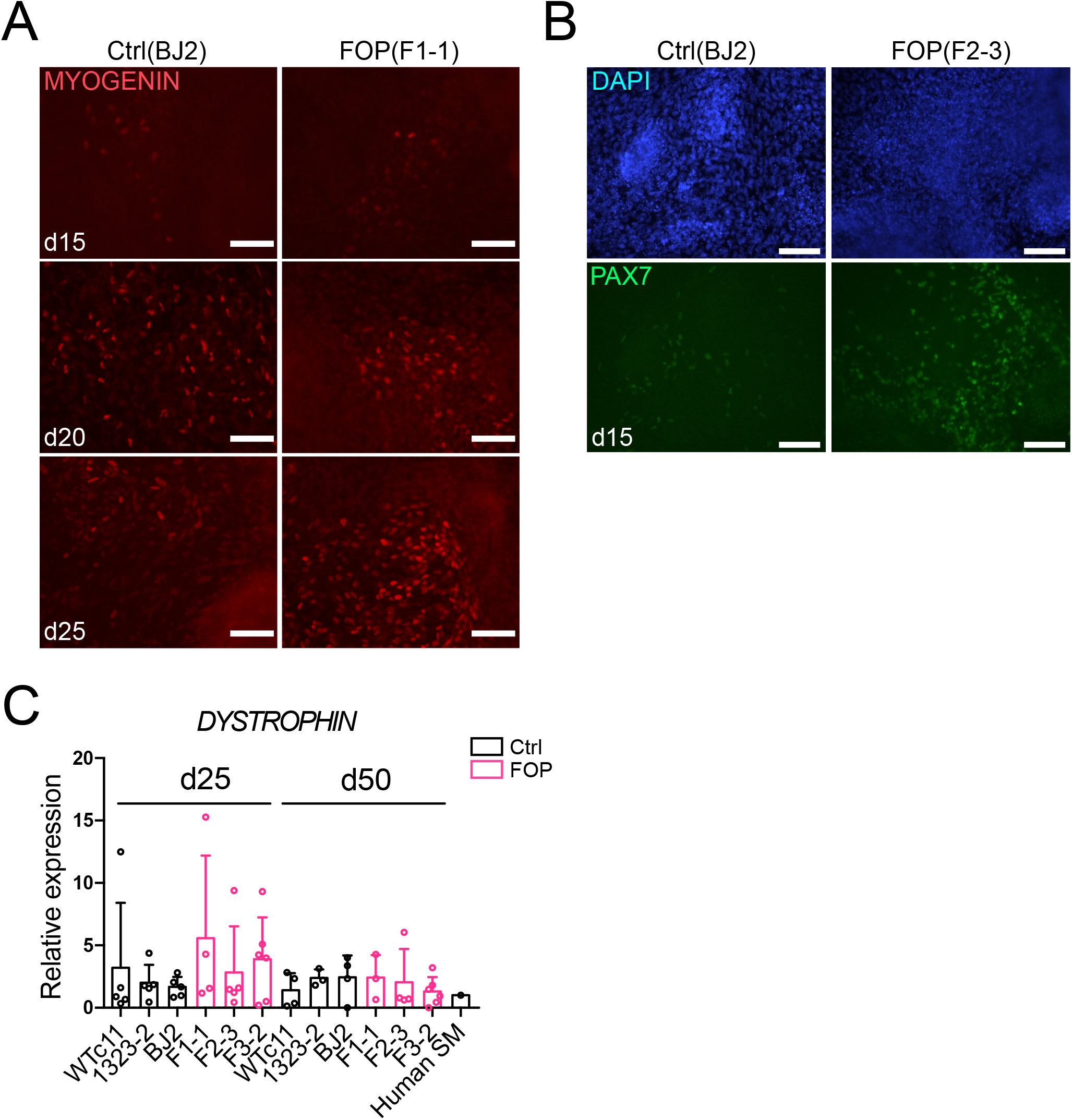
Early expression of muscle markers during the myogenic differentiation. **(A)** MYOGENIN (scale bar, 200µm) and **(B)** PAX7 immunofluorescence staining showing that MYOGENIN and PAX7 are expressed as early as day 25 during the myogenic differentiation in control and BMP impaired hiPSCs (scale bar, 200µm). **(C)** *DYSTROPHIN* gene expression at day 25 and day 50 of differentiation (n=3 biological replicates and n≥3 technical replicates). Error bar represent mean and SD.

**Figure 3-figure supplement 1.**
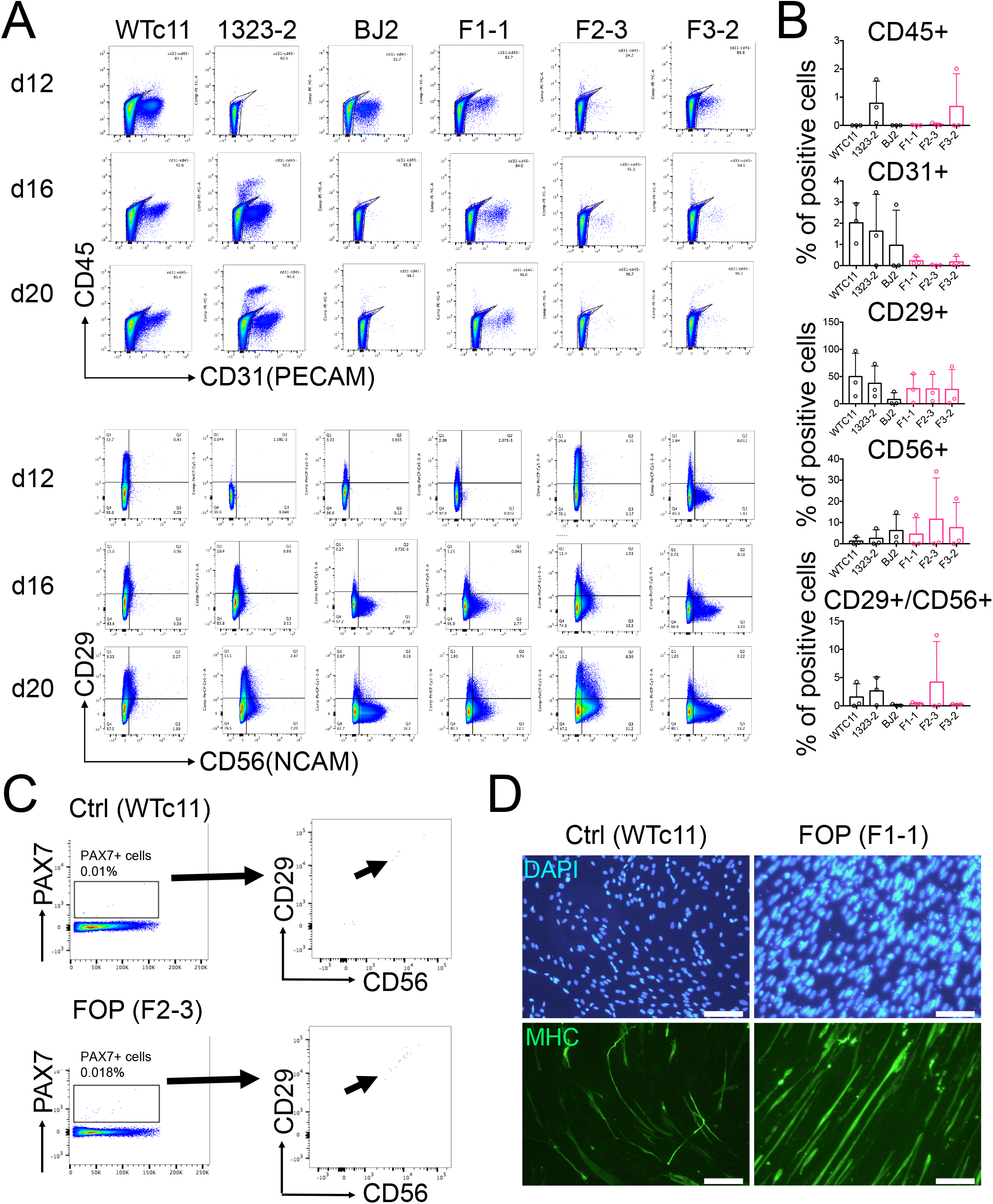
Differentiated cells types in the myogenic differentiation varies between cell lines, sorted cells express PAX7 and can differentiate into myotubes. Related to Figure 3. **(A)** Representative flow cytometry profiles of negative (CD45 and CD31) and positive (CD29 and CD56) markers used to purify muscle stem cells. **(B)** Quantification of the percentage of cells positive for CD45, CD31, CD29, and CD56 at day 20. (n=3 biological replicates and n=3 technical replicates). Error bar represent mean and SD. **(C)** FACS analysis of the myogenic differentiation at day 50. Muscle stem cells stained for PAX7, CD29, and CD56. **(D)** Representative immunofluorescence staining for MHC showing sorted muscle stem cells after being cultured in differentiation media for 7 days (scale bar, 200µm).

**Figure 4-figure supplement 1.**
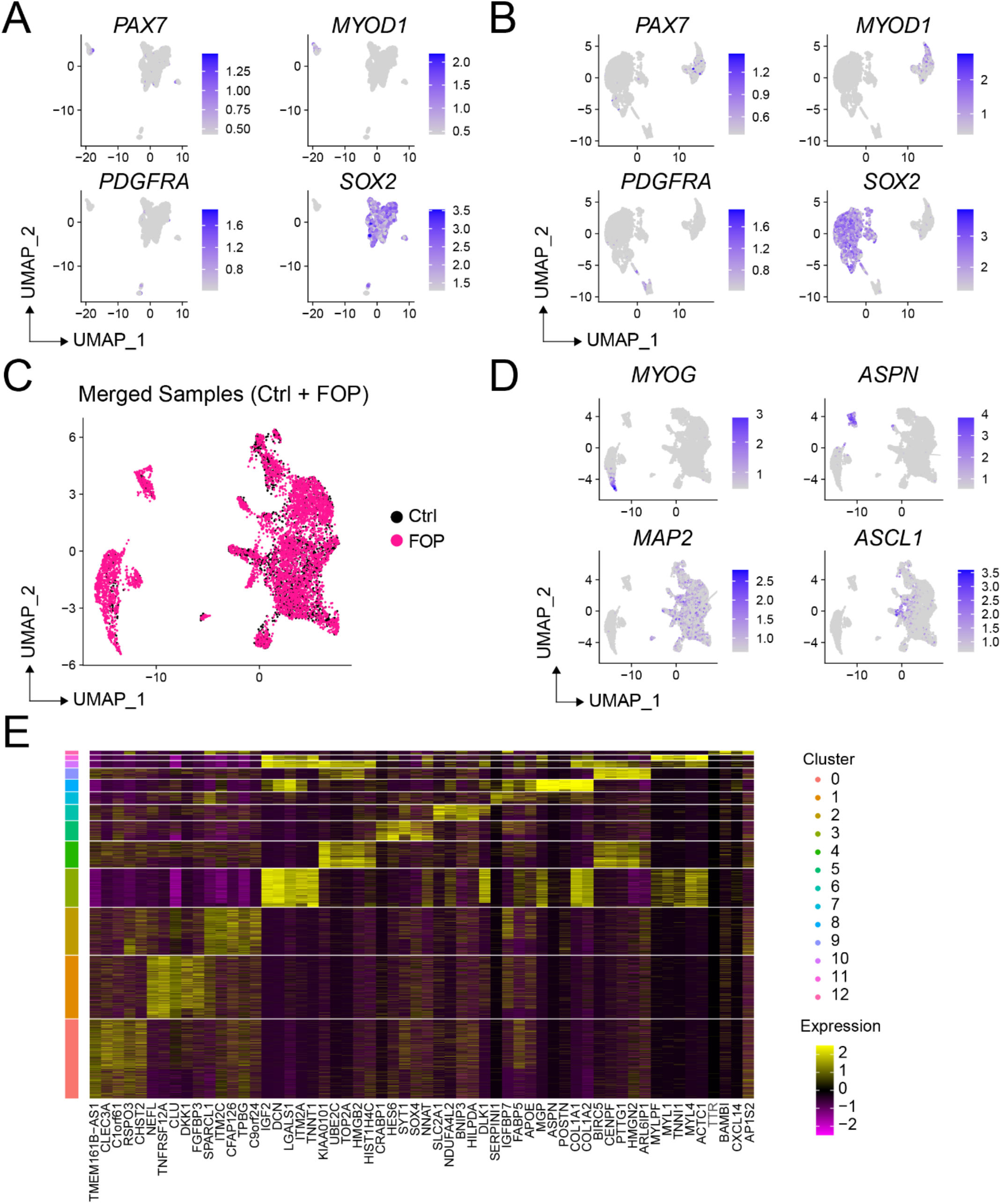
Markers expressed in sorted control and FOP HNK1^-^CD45^-^CD31^-^ CXCR4^+^CD29^+^CD56^dim^ cells. **(A-B)** Feature expression plots showing the localization of cells expressing myogenic markers (*PAX7, MYF5*) mesenchymal marker (*PDGFRA*) and neuronal progenitor marker (*SOX2*) in the control (A) and FOP **(B)** sorted cells. **(C)** UMAP visualization plots of merged samples with cells colored by samples. **(D)** Feature expression plots showing the localization of cells expressing myogenic (MYOG), mesenchymal (*ASPN*) and neuronal markers (*MAP2, ASCL1*) in the merged samples. **(E)** Heat map of the top 5 differentially expressed genes in each cluster of the merged sample analysis.

**Figure 5-figure supplement 1.**
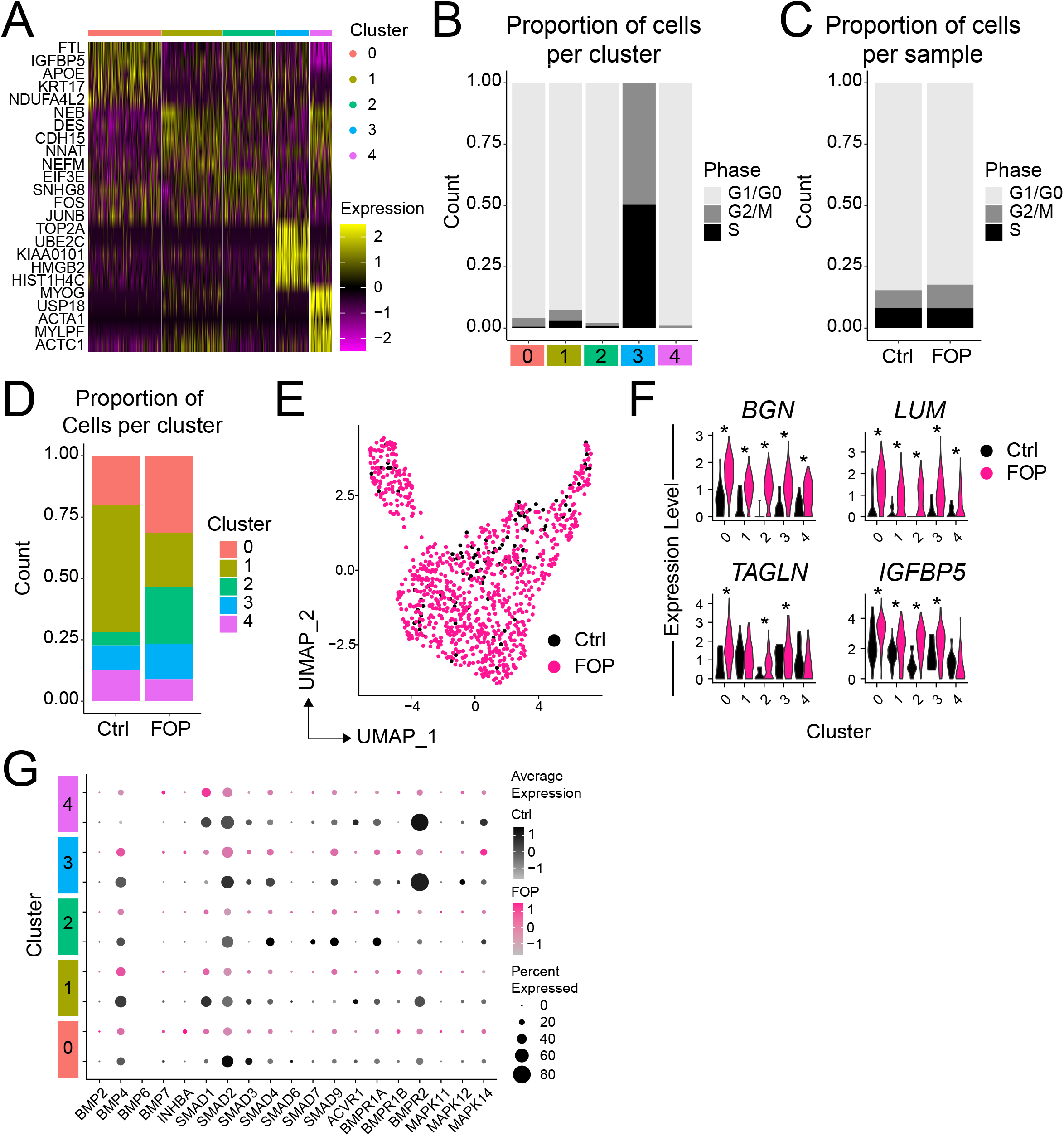
Analysis of the myogenic sub-cluster. **(A)** Heat map of the top 5 differentially expressed genes in each cluster. **(B)** Cycle genes were scored for each cluster. Bar plot depicting the proportion of cells in G1/G0, G/2M and S phase for each cluster. **(C)** Proportion of cells in G1/G0, G2/M and S phase for each sample. **(D)** Proportion bar graph of cells per cluster for the control and FOP samples. **(E)** UMAP showing the distribution of the two merged samples. **(F)** Additional genes significantly differentially expressed by more than 1-fold. (**G)** Dot plot displaying the expression of genes associated with BMP and P38MAPK pathways for each cluster.

**Figure 6-figure supplement 1.**
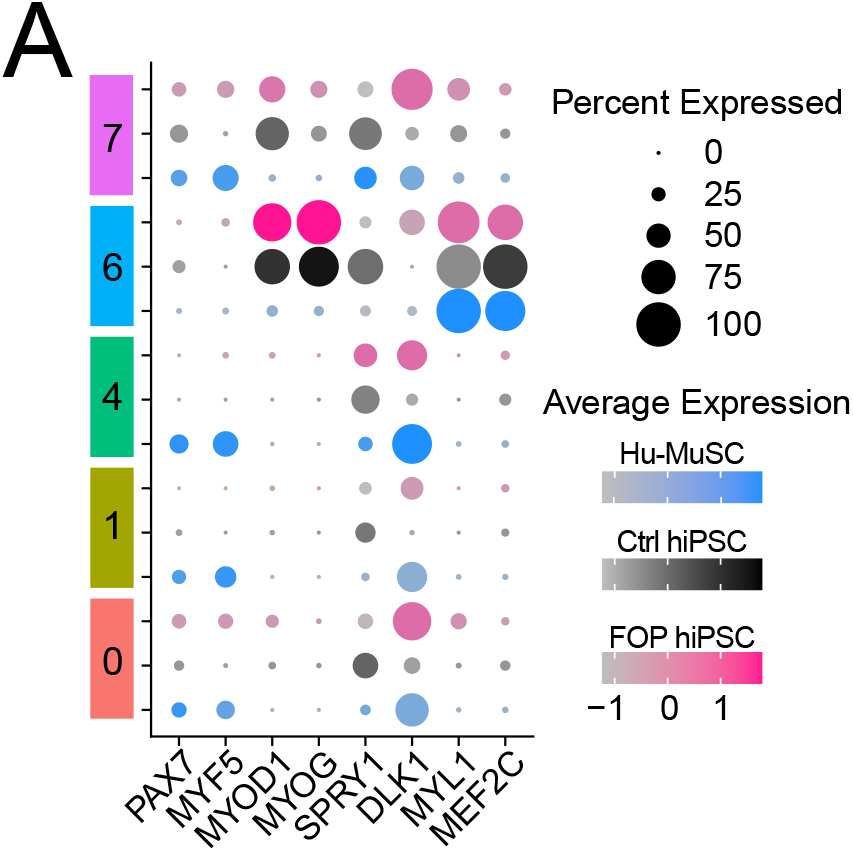
Expression of myogenic markers across primary and hiPS-derived myogenic cells. Dot plot displaying the average expression and the percent of cells expressing myogenic genes across clusters for the 3 merged samples.

**Figure 1-Source Data 1.**
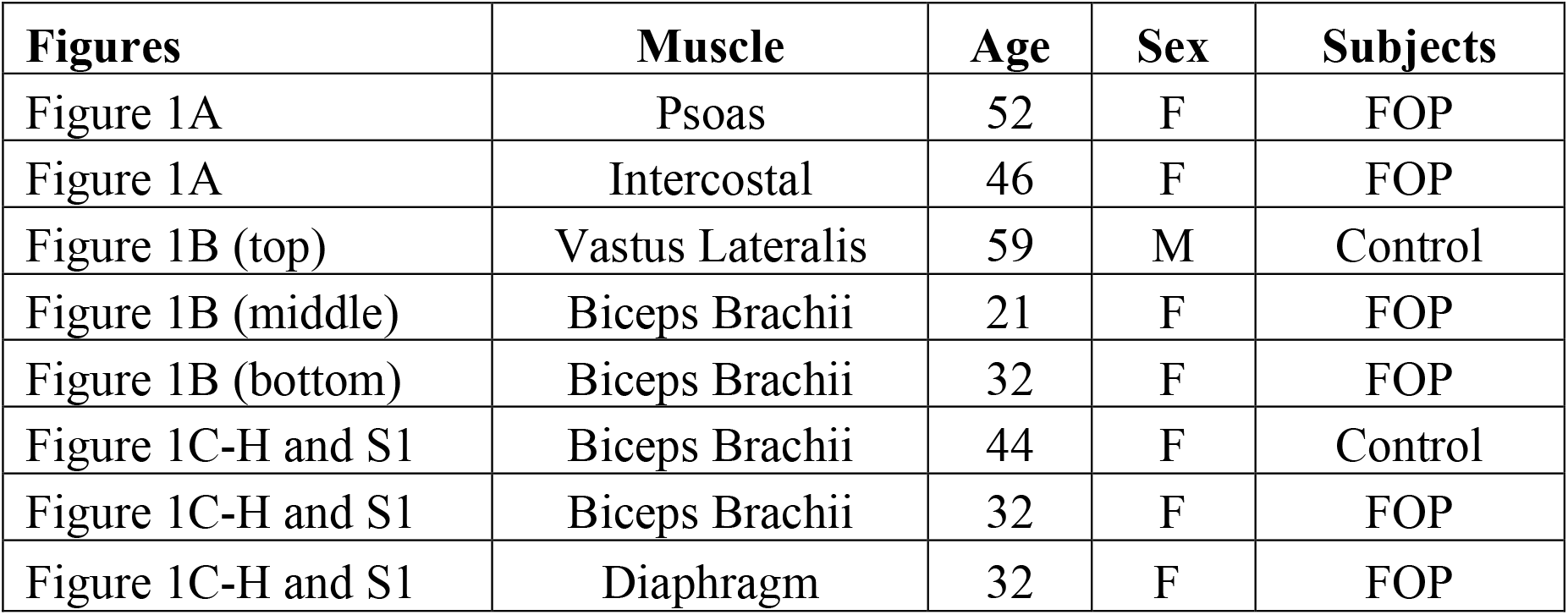
Muscle specimen information.

**Figure 1-Source Data 2.**
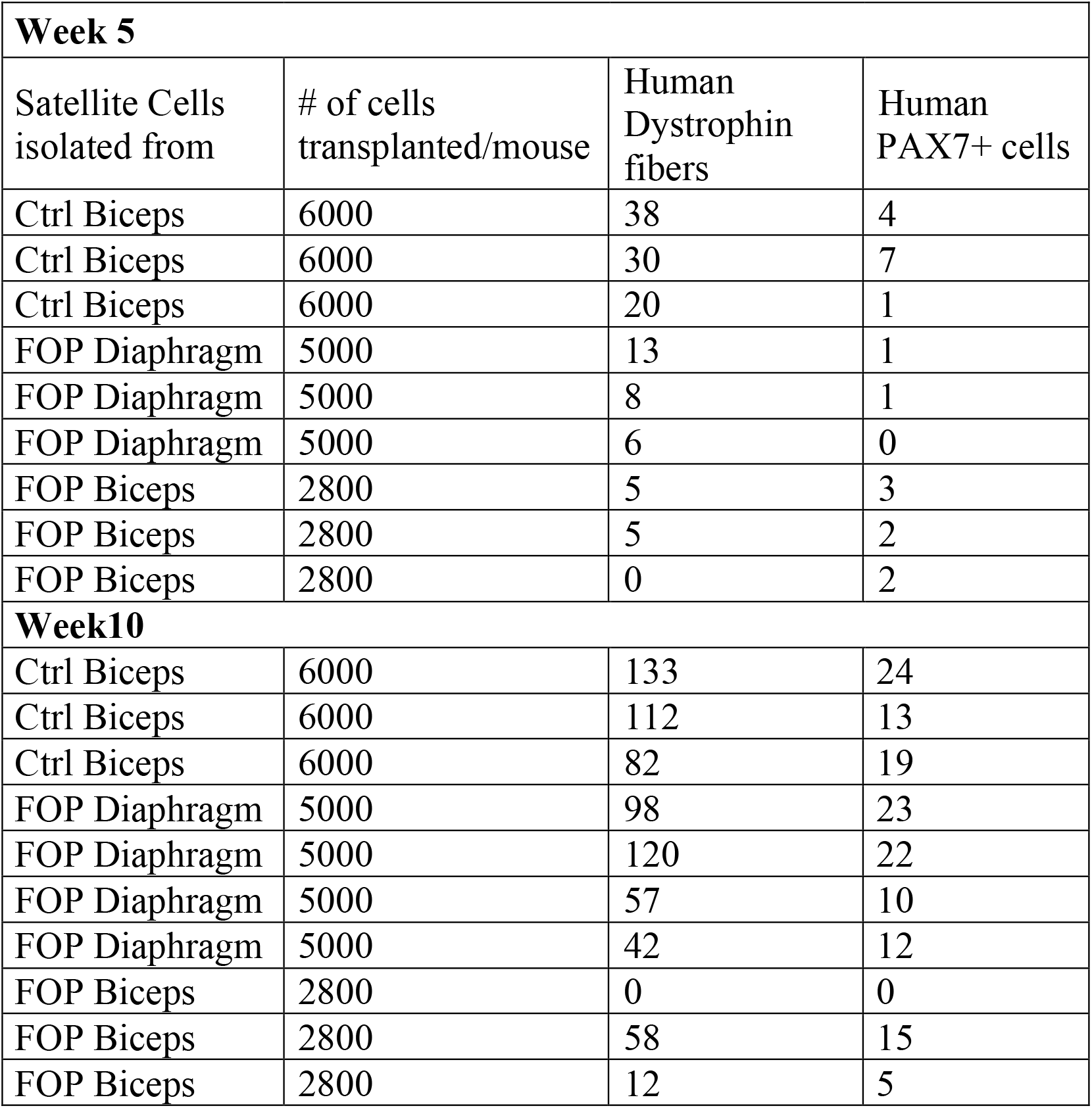
Hu-MuSCs transplantation details.

**Figure 3-Source Data 1:**
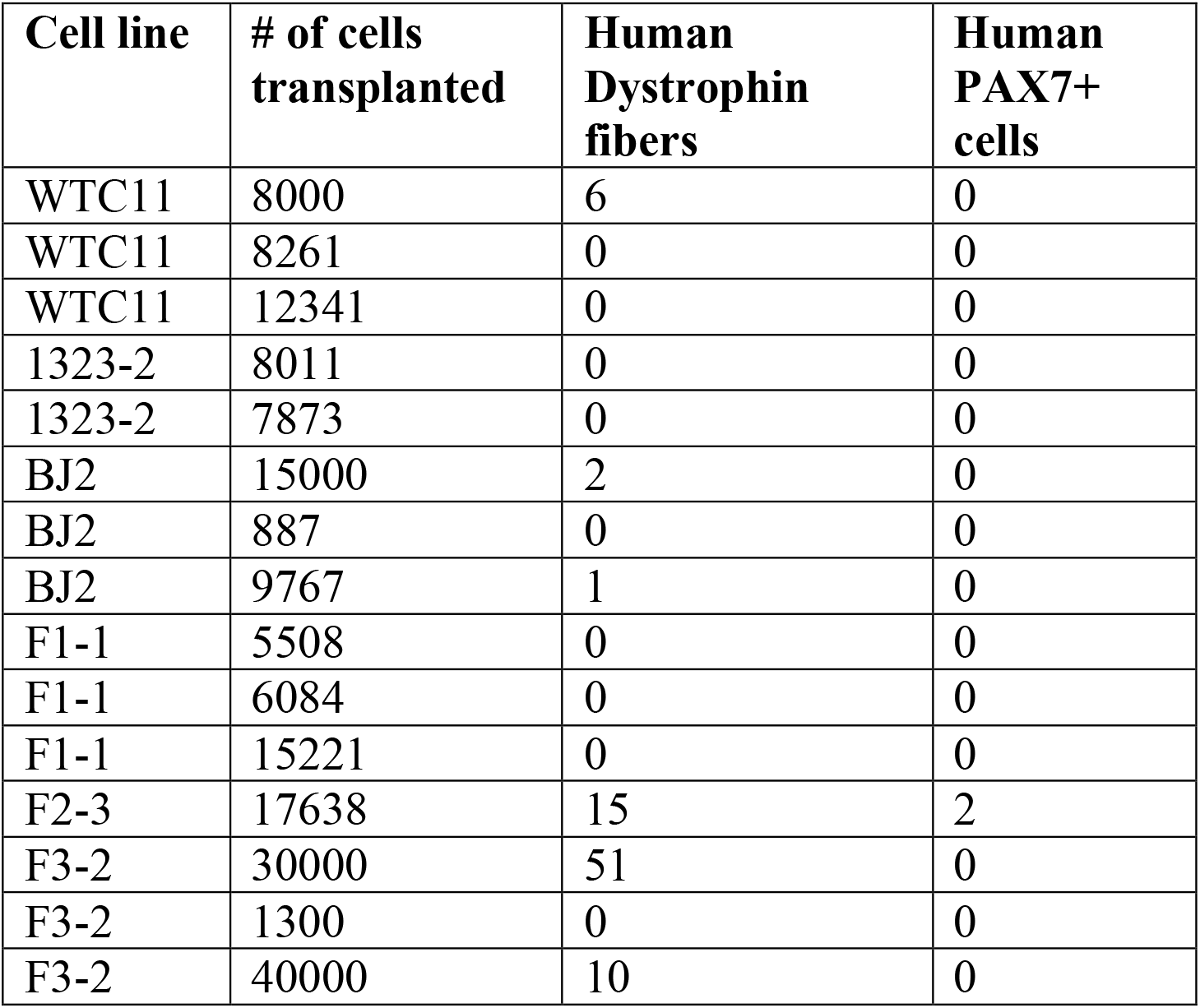
hiPSC-derived HNK1^-^CD45^-^CD31^-^ CXCR4^+^CD29^+^CD56^dim^ cell transplants.

**Figure 4-Source Data 1:**
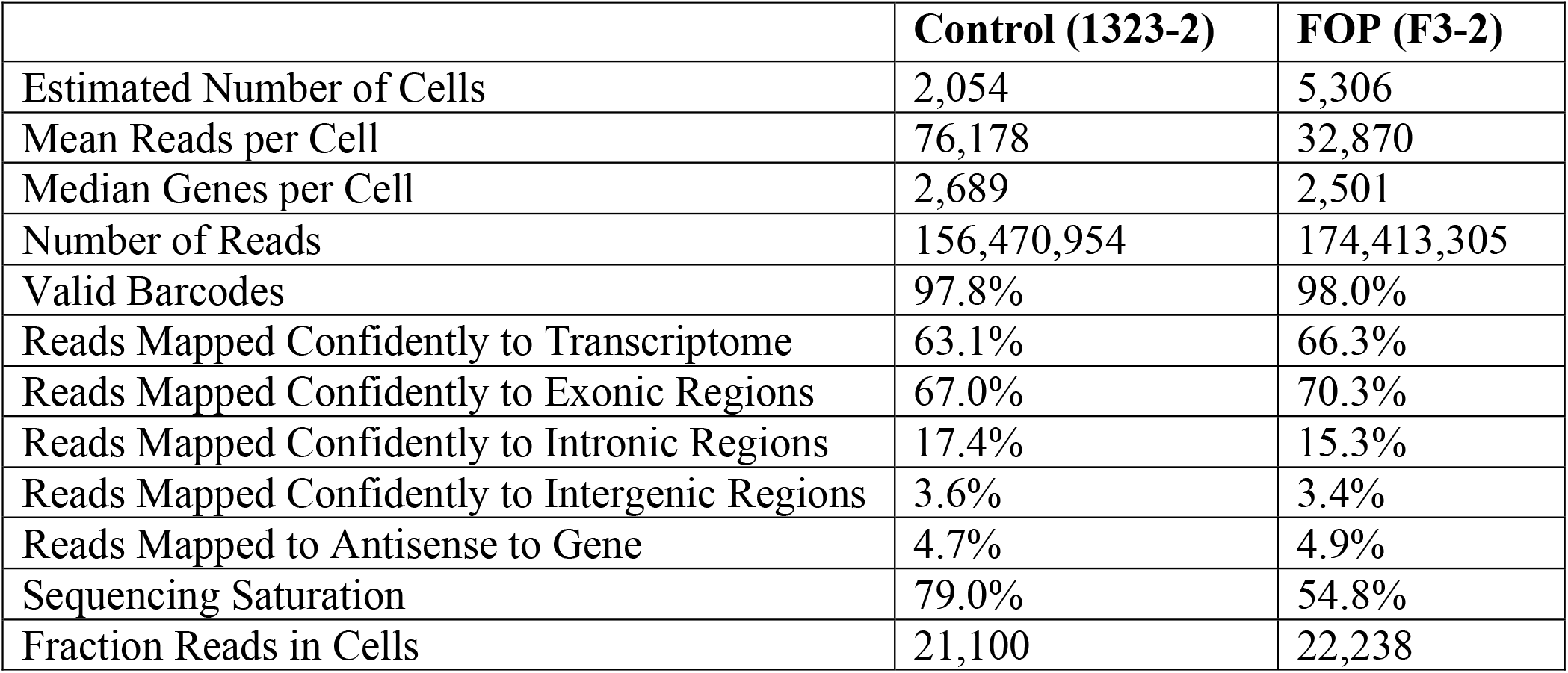
Quality control information for each sample.

**Figure 5-Source Data 1. Differential expression analysis of control and FOP myogenic cells (see excel file).**

**Figure 6-Source Data 1. BEAM analysis (see excel file).**

## SUPPLEMENTAL EXPERIMENTAL PROCEDURES

### Flow cytometry

Cells were treated with Accutase for 20min at 37C, washed with FACS buffer, and stained for HNK1-PE, CD45-PE, CD31-PE, CD56-APC, CXCR4-PE-Cy7, and CD29-FITC (eBioscience). HNK1^-^CD45^-^CD31^-^ cells [to select against neuronal cells (HNK1, Human Natural Killer-1) (Choi et al., 2016; Hicks et al., 2018), hematopoietic cells (CD45), and endothelial cells (CD31)], co-expressing CD29, CXCR4, and intermediate CD56, markers present on human PAX7^+^ cells (Garcia et al., 2018; Garcia et al., 2017; Xu et al., 2015) were sorted with a FACSAriaIII (BD Biosciences) and Sytox Blue (Life Technologies) was used as a viability marker. Alternatively, cells were permeabilized and fixed (Fix/Perm Buffer Set, BioLegend). Fixed cells were first incubated with primary antibodies (PAX7 (DSHB), CD29 (BD Biosciences) and CD56 (BD Biosciences) following by secondary antibodies (Alexa350-conjugated goat anti-mouse IgG, Alexa488-conjugated goat anti-rat IgG and Alexa546-conjugated donkey anti-goat IgG, Life Technologies).

### Cell Immunostaining and NSG Tibialis Anterior Analysis

Collected tibialis anterior frozen cross sections were fixed in 4% PFA for 10 min at room temperature, washed with PBST (PBS with 0.1% Tween-20 (Sigma-Aldrich) and blocked in PBS with 10% goat serum for 1hr at room temperature. Slides were then incubated 4hrs at room temperature with the following: mouse monoclonal anti-human DYSTROPHIN (DSHB), mouse monoclonal IgG1 anti-PAX7 (DSHB), rabbit polyclonal anti-Laminin (Sigma-Aldrich), mouse monoclonal IgG2b anti-human SPECTRIN (Leica Microsystems), and mouse monoclonal IgG2b anti-human LAMIN A/C (Vector Laboratories). After PBST wash, slides were incubated with the following secondary antibodies: Alexa Fluor 555 goat anti-mouse IgG, Alexa Fluor 594 goat anti-mouse IgG1, Alexa Fluor 488 goat anti-mouse IgG2b, and Alexa Fluor 647 goat anti-rabbit (Life Technologies). Finally, sections were mounted with VECTASHIELD mounting media with DAPI (Vector Laboratories). All samples were examined using a Leica upright or DMi8 Leica microscope. Sections with the most human fibers were used for human DYSTROPHIN and PAX7 quantification for each condition.

### Single cell RNA Sequencing and Analysis

45,000 (FOP) and 30,000 (control) HNK1^-^CD45^-^CD31^-^ CXCR4^+^CD29^+^CD56^dim^ cells isolated from the iMPC differentiations were loaded onto one well of a 10X chip to produce Gel Bead-in-Emulsions (GEMs). GEMs underwent reverse transcription to barcode RNA before cleanup and cDNA amplification. Libraries were prepared with the Chromium Single Cell 3’ Reagent Version 2 Kit. Each sample was sequenced on 1 lane of the NovaSeq 6000 S4. Sequencing reads were processed with Cell Ranger version 2.0.0. using the human reference transcriptome GRCh38. The estimated number of cells, mean reads per cell, median genes per cells, median UMI (Unique Molecular Identifier) counts per cells as well as other quality control information are summarized in **Figure 4-Source Data 1**. Gene-barcoded matrices were analyzed with the R package Seurat v3.1.5 (Satija, Farrell, Gennert, Schier, & Regev, 2015; Stuart et al., 2019; Team, 2014; Zheng et al., 2017). Gene core matrices from single cell RNA sequencing of primary human satellite cells isolated from a vastus muscle (Barruet et al., 2020) was used when comparing the transcriptional profile of hiPS-derived HNK1^-^CD45^-^CD31^-^ CXCR4^+^CD29^+^CD56^dim^ cells. For the comparison with primary Hu-MuSCs, hiPSC-derived cell sequencing reads were re-aligned using the human reference transcriptome hg19. Cells with fewer than 500 genes, greater than 5000 genes and genes expressed in fewer than 5 cells were not included in the downstream analyses. Cells with more than 10% mitochondrial counts were filtered out. Samples were normalized with NormalizeData using default settings. The FindVariableFeatures function was used to determine subset of feature that exhibit high cell-to-cell variation in each dataset based on a variance stabilizing transformation (“vst”). We used the default setting returning 2,000 feature per dataset. These were used for downstream analysis. In the case of the merged data analysis samples were combined utilizing the FindIntegrationAnchors function with the ‘dimensionality’ set at 30. Then, we ran these “anchors” to the IntegratData function for batch correction for all cells enabling them to be jointly analyzed. The resulting outputs were scaled mitochondrial contamination regressed out with the ScaleData function. In addition, while we didn’ t regress out heterogeneity associated with cell cycle stage since it is an important factor in determining the state of quiescence of our sorted human muscle stem cells, we regressed out differences between G2/M and S cell cycle stage. PCA was performed with RunPCA, and significant PCs determined based on the Scree plot utilizing the function PCElbowPlot. The resolution parameter in FindClusters was adjusted to 0.5. Clusters were visualized by UMAP with Seurat’ s RunUMAP function. We performed differential gene-expression utilizing Seurat v3’ s FindMarkers function with default settings which utilizes the Wilcoxon rank-sum test to calculate adjusted p values for multiple comparisons. We used the CellCycleScoring function to assign score based on the expression of G2/M and S phase markers (Regev et al., 2017). Myogenic cells were further analyzed by sub-clustering using the subset function. We then use the FindNeighbors (dims=15) and FindClusters (resolution = 0.4) functions on the myogenic cell subset of the merged hiPSCs samples only to identify sub-clusters corresponding to different myogenic states. To order the cells in pseudotime based on their transcriptional similarity we used Monocle 2.12. Variable genes from Seurat analysis were used as input and clusters were projected onto the minimum spanning tree after ordering. Gene expression patterns were plotted with plot_genes_branched_heatmap, plot_genes_branched_pseudotime, and plot_multiple_branches_pseudotime. The BEAM (branch expression analysis modeling) function was used to score gene significance in a branch-dependent manner. Cells were re-ordered using the orderCells function to set branch A (myocytes) in **Figure 5F** as the “root-state”. This allowed us to determine genes that were significantly branch dependent in branch B (mainly Hu-MuSCs) vs branch C (hiPS-derived cells) using the BEAM analysis.

